# Local sequence context determines the effect of glycine substitutions in collagen triple helices

**DOI:** 10.64898/2026.06.01.729402

**Authors:** Anton V. Persikov

## Abstract

Glycine substitutions in the collagen triple helix cause diverse heritable disorders, but their effects vary with local sequence environment. We tested whether sequence context helps determine the consequences of Gly replacement by combining case-weighted bioinformatic analysis, thermodynamic measurements on collagen model peptides (CMPs), and all-atom molecular dynamics simulations. Case-weighted analysis of pathogenic *COL3A1* glycine substitutions identified Pro immediately following the substituted Gly, corresponding to a GP context, as the strongest enriched local feature. To examine this experimentally, we designed CMPs with stabilizing terminal segments flanking native collagen sequence windows containing clinically observed Gly→Ser and Gly→Arg substitutions. Gly→Arg substitutions were generally more destabilizing than Gly→Ser at the same site. However, the strongest effect was sequence-dependent: within the Gly→Ser class, GP-site substitutions caused larger losses of thermal stability and unfolding enthalpy than nonGP substitutions, and some GP-site Gly→Ser mutations were as destabilizing as Gly→Arg substitutions. Molecular dynamics simulations showed that all peptides remained globally triple-helical, but GP-site mutants exhibited greater loss of canonical interchain hydrogen bonds and reduced local backbone accommodation. Thus, the effect of glycine substitution in collagen depends not only on the replacing residue but also on the permissiveness of the surrounding sequence, with Pro-adjacent sites representing especially restrictive local environments.

**Statement of Significance:** Collagen diseases often result from replacement of a required glycine in the triple helix, but the same substitution can have very different consequences at different sites. Using clinical bioinformatics, collagen model peptides, and molecular dynamics simulations, we show that the surrounding sequence is a major determinant of mutational outcome. The most restrictive context identified here is a GP site, in which proline immediately follows the substituted glycine: Gly→Ser mutations at GP sites are more destabilizing than Gly→Ser mutations at sites without a following proline and can be as damaging as Gly→Arg substitutions. These results define a sequence-based rule for collagen mutational tolerance relevant to both triple-helix stability and interpretation of pathogenic variants.

## Introduction

Collagens are defined by a deceptively simple structural rule: a long, continuous triple helix formed by three polyproline-II-like chains in which glycine occupies every third position. These long triple-helical molecules constitute a key structural element of animal extracellular matrices, assembling into fibrils and networks that provide tissues with tensile strength while also presenting recognition sites for cells and soluble ligands (1). This balance between mechanical integrity and biological function becomes especially clear in the “collagenopathies,” where single-amino-acid changes can derail folding, secretion, assembly, and tissue function. Classic examples include osteogenesis imperfecta (OI), most often caused by variants in *COL1A1* and *COL1A2*, which encode the α1(I) and α2(I) chains of type I collagen, respectively (2,3), and vascular Ehlers-Danlos syndrome (vEDS), caused by mutations in *COL3A1*, which encodes the α1(III) chain of type III collagen (4,5).

A striking feature of the collagen triple helix is how narrowly its thermodynamic stability is tuned. Across species, collagen unfolding temperatures track the temperatures at which organisms live, spanning polar ectotherms to hydrothermal-vent fauna, consistent with evolutionary tuning of triple-helix stability (6). At the same time, careful equilibrium measurements showed that the melting temperature of human type I collagen monomers in physiological solution lies several degrees *below* 37 °C, implying that collagen is not designed to be maximally stable as an isolated molecule (7,8). This paradox becomes less surprising in the broader context of collagen biology: collagens fold *in vivo* in a crowded, propeptide- and chaperone-assisted intracellular environment and then become further stabilized by the supramolecular packing of fibrils and networks (9). Moreover, the functional collagen molecule cannot behave as an inert crystalline rod, because enzymatic remodeling, regulated proteolysis, and reversible ligand binding all require local fluctuations and transient accessibility. In this view, “fine-tuning” of stability is not a subtlety but a design constraint: the triple helix must be stable enough to fold and persist, yet labile enough to participate in assembly and interactions on biological timescales.

Consistent with this, collagen stability is not uniform along the sequence. High-resolution structures of collagen-like peptides revealed that local amino-acid content produces measurable variations in twist, hydration, and packing within a globally conserved fold (1,10). Structural heterogeneity is also evident at larger scales, where the D-periodic organization of fibrils imposes a repeating landscape of molecular environments and accessibility (11). More recently, the concept of alternating rigid and flexible zones has been formalized for collagen III as a “flexi-rod” architecture, in which flexible sequences are interspersed with bioactive domains linked to hemostatic interactions (12). Together, these observations argue against treating collagen as a mechanically and thermodynamically uniform polymer. Instead, sequences can encode local differences in stability and mechanics, which in turn can regulate when and where functional sites are exposed (10,12,13).

The use of short self-associating synthetic collagen model peptides (CMPs) has been central to establishing how sequence encodes triple-helix structure and stability. Early crystallographic work on CMPs provided an atomic description of the collagen fold and a framework for interpreting sequence-dependent deviations from an “ideal” triple helix (10,14). Systematic host-guest CMP thermodynamics allowed determination of residue propensity scales including all Gly, Xaa, and Yaa repeating positions (15,16). These studies also identified specific interaction motifs as major stability determinants, including strong contributions from electrostatic networks and favorable Lys-containing charge patterns such as KGE and KGD (17). Beyond homotrimers, CMP design has expanded toward controlled heterotrimer formation and higher-order self-assembly, notably through electrostatic programming and “sticky-ended” architectures that connect triple helices into larger materials (18–20). However, modeling pathological glycine substitutions in short CMPs remains challenging, because glycine replacements can strongly destabilize the fold, produce partially folded intermediates, and require stabilizing agents (16).

Here, we tested the hypothesis that the local amino acid environment surrounding a glycine substitution is a major determinant of the magnitude of triple-helix destabilization caused by that mutation. To do so, we used a modified CMP design that builds on the stabilizing-host concept but differs from classical host-guest peptides by placing a relatively long native collagen-derived sequence window between terminal stabilizing segments. This design maintains a well-folded triple-helical scaffold while preserving the native local sequence context for testing single Gly substitutions under standard solution conditions without stabilizing agents. We combined thermodynamic measurements on mutation-bearing peptides, case-weighted bioinformatic analysis of residues flanking clinical Gly substitutions, and all-atom molecular dynamics simulations to examine whether local sequence context helps explain why the same glycine substitution can have markedly different structural consequences in different collagen regions.

## Materials and Methods

### Sequence-context enrichment analysis of case-weighted Gly substitutions

To test whether specific residues are enriched at defined positions flanking pathogenic glycine substitutions in the homotrimeric collagen triple helix, pathogenic *COL3A1* variants associated with vascular Ehlers-Danlos syndrome (vEDS) were assembled from the public Leiden Open Variation Database (LOVD) (21). Only pathogenic variants were retained. Duplicate published records were handled conservatively: repeated patient-level entries within a source were preserved, whereas cross-source redundancies were removed only when the same variant was linked to the same publication reference. After restriction to glycine missense substitutions within the uninterrupted triple-helical region, the final case-weighted dataset contained 744 missense cases at 343 unique Gly sites.

For each substituted Gly site, the local triple-helical register was defined by assigning the mutated residue as the central Gly and extracting residues at eight flanking positions: *X_-2_, Y_-2_, X_-1_, Y_-1_,* (two triplets upstream) and *X_1_, Y_1_, X_2_, Y_2_* (two triplets downstream). Flanking residues were obtained directly from the α1(III) chain sequence (UniProt ID: P02461). A site-specific case weight was defined as the total number of reported glycine missense substitutions at that site.

Expected background frequencies were determined by enumerating all Gly positions in the main triple-helical region of α1(III) chain for which each flanking position could be assigned unambiguously, excluding sites truncated by sequence boundaries. For each flanking position *i* and amino acid *a*, the background probability *f_i_(a)* was calculated as the fraction of eligible reference Gly sites with residue *a* at position *i*. The observed case-weighted count *n_i_(a)* was calculated as the sum of case weights over all mutated Gly sites with residue *a* at position *i*. The total number of case-weighted observations contributing to position *i* was *N_i_*. Enrichment was tested with a one-sided binomial test under the null hypothesis that the frequency of residue *a* at position *i* among mutation cases matches the corresponding background probability *f_i_(a)*. Under the null, *n_i_(a)* follows a binomial distribution, *X_i_(a)* ∼ *Binomial(N_i_, f_i_(a))* with parameters *N_i_* and *f_i_(a)*, and *p*-values were computed as the upper-tail probability *P(X_i_(a) ≥ n_i_(a))*.

Because 20 amino acids were tested at eight flanking positions (160 hypotheses), *p*-values were adjusted across all tests by the Benjamini-Hochberg false discovery rate procedure (22). Adjusted *p*-values are reported as *q*-values. The fold-enrichment was computed as *n_i_(a)/E_i_(a)*, where the expected count under the null *E_i_(a) = N_i_ f_i_(a)*. Results were visualized as a volcano plot with log_2_(fold-enrichment) on the x-axis and -log_10_(*q-*value) on the y-axis. Residue-position combinations with q < 0.05 were considered significantly enriched and were annotated by position and amino acid identity.

### Collagen model peptides

Short triple-helical peptide systems have been described as “collagen model peptides”, “collagen-like peptides”, and “collagen-mimetic peptides” (18,23,24). Here we use “collagen model peptides” (CMPs) to emphasize their role as controlled peptide models of native collagen sequences. CMPs were designed to examine the effects of pathogenic glycine substitutions observed in osteogenesis imperfecta (OI) and vascular Ehlers-Danlos syndrome (vEDS) within native local sequence environments while retaining sufficient triple-helical stability for biophysical analysis. In peptide sequences, Hyp denotes 4(R)-hydroxyproline, which is represented as O in the one-letter notation used here. Each peptide contained 38 residues, with the mutation site centered within a four-triplet collagen-derived sequence window flanked by stabilizing (Gly-Pro-Hyp)_4_ segments at both ends. Matched WT and mutant peptides carrying Gly→Ser (G→S) and/or Gly→Arg (G→R) substitutions differed only at the central Gly position (Figure 1).

**Figure 1.**
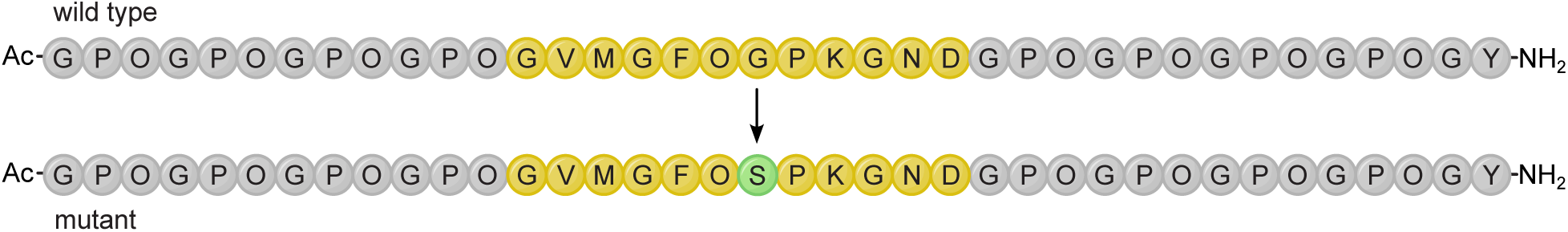
Design of collagen model peptides for site-specific analysis of pathogenic glycine substitutions. Each peptide contains 12 Gly-Xaa-Yaa triplets per chain, with stabilizing (GPO)_4_ segments at the N- and C-termini (grey shading) flanking a central four-triplet segment derived from real collagen sequence (yellow shading). Wild-type and mutant CMPs were designed by introducing disease-associated Gly→Ser and/or Gly→Arg substitutions (green shading) within the native central sequence context. The repeating sequence began and ended with Gly, and a C-terminal Tyr was included for concentration determination. The N terminus was acetylated and the C terminus amidated to reduce end effects and minimize intermolecular association. The sequence example represents T3-G415 and T3-G415S peptides.

CMPs were named according to collagen chain and the position of the substituted Gly residue within the main Gly-Xaa-Yaa repeating domain. Thus, in A1-G382S, “A1” denotes the α1(I) chain, “A2” for the α2(I) chain, and “T3” denotes the α1(III) chain; G382S denotes substitution of Gly382 by Ser.

Peptides were synthesized by solid-phase methods at the Tufts University Core Facility (Boston, MA). Crude products were purified to >95% on a C18 column using a Shimadzu reverse-phase HPLC system with 0.1% trifluoroacetic acid and a 0-40% acetonitrile/water gradient. Peptide identity was confirmed by matrix-assisted laser desorption/ionization mass spectrometry (MALDI-MS). Summary characterization data are provided in the Supplemental Material as Table S1. The complete set of RP-HPLC chromatograms and MALDI-TOF mass spectra for all 27 synthesized peptides is available in the public Zenodo repository (DOI: 10.5281/zenodo.20560423). Purified peptides were dissolved in phosphate-buffered saline containing 0.15 M NaCl and 10 mM sodium phosphate, pH 7.0. Peptide concentrations were typically 1-2 mg/mL and were determined from Tyr absorbance at 275 nm using ε^275^ = 1400 M^−1^ cm^−1^. Before biophysical measurements, peptide solutions were annealed by heating to approximately 75 °C for 10-15 min in sealed tubes, cooling slowly to near the expected peptide Tm with incubation for 3-4 h, incubating at 25 °C for 3-4 h, and storing overnight at 5 °C.

### Circular dichroism spectroscopy

CD measurements were made on an Aviv Model 62DS spectrometer. Peptide solutions were studied in 1-mm quartz cuvettes using a rotating four-cell holder. For thermal melting experiments, ellipticity at 225 nm was monitored as temperature was increased from 0 to 90 °C at an average heating rate of 0.1 °C/min under standard conditions. As in our earlier work, these heating rates do not produce equilibrium melting curves, but the transitions approximate a two-state model and the apparent *T_m_* values are reproducible and useful for comparison among peptides (25,26). A trimer-monomer two-state model was used to analyze the thermal transitions.

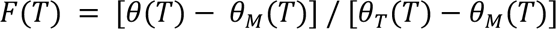

where *θ* is the observed ellipticity and *θ_T_* and *θ_M_* are the ellipticities in the native and monomer states, respectively, each treated as a linear function of temperature. *T_m_* values were taken as the midpoint of the transition (*F* = 0.5) under these defined heating conditions. From independently prepared samples, the error in *T_m_* determination was < 0.4 °C.

### Differential scanning calorimetry

DSC experiments were performed on a Nano-DSC II instrument (Calorimetry Sciences Corp., Model 6100) at a scan rate of 1.0 °C/min from 0 to 90 °C. Baseline correction of buffer-subtracted scans was performed by fitting smoothing splines to the pre-transition and post-transition regions, followed by linear interpolation across the transition. The excess heat capacity curve was obtained after baseline subtraction, and the transition enthalpy (*ΔH*) was calculated by numerical integration of the excess heat capacity peak over the temperature range spanning the transition. The experimental error in calorimetric enthalpy determination did not exceed 5%.

### Statistical analysis of biophysical comparisons

For each mutant peptide, destabilization relative to the matched WT control was expressed as *ΔT_m_ = T_m_(WT) - T_m_(mutant)*, using CD-derived *T_m_* values, and as *ΔΔH = ΔH(WT) - ΔH(mutant)*, using DSC-derived calorimetric enthalpy values. Group comparisons were performed at the peptide level using two-tailed Mann-Whitney U tests.

### Molecular dynamics simulations

All-atom molecular dynamics (MD) simulations were performed on WT and mutant CMP triple helices in explicit CHARMM TIP3P water using OpenMM (27) at 300 K. Initial systems were prepared with CHARMM-GUI using N-terminal acetylation and C-terminal amidation to match the experimental peptide design (28). Each peptide was placed in a periodic water box with at least 5 Å padding from the solute, giving final box dimensions of approximately 125 × 125 × 125 Å. Systems were neutralized and supplemented with 0.15 M NaCl to match the experimental buffer conditions, resulting in approximately 174 Na^+^ and 174 Cl^−^ ions per system. The CHARMM-GUI-generated hydrated input systems for all simulated CMPs, including PDB, CRD, PSF, and system information files, have been deposited in Zenodo at doi:10.5281/zenodo.20750355.

Each CMP was simulated in a 500 ns production run using the CHARMM36mGP force field, a modification of CHARMM36m in which the CMAP correction for Gly, Pro, and Hyp backbone dihedrals is scaled by 0.5 to improve simulation of collagen triple helices (29). Long-range electrostatics were treated with particle mesh Ewald using a 1.0 nm real-space cutoff and an Ewald error tolerance of 0.0005. Bonds involving hydrogen were constrained, water molecules were rigid, and hydrogen mass repartitioning to 2.0 amu permitted a 4 fs integration time step. Simulations were performed in the NPT ensemble at 300 K and 1 atm using a Langevin middle integrator with a friction coefficient of 1.0 ps^-1^ and a Monte Carlo barostat applied every 25 steps. After energy minimization and brief equilibration following velocity assignment at 300 K, production trajectories were propagated for 125,000,000 steps per run. Simulations were run on the CUDA platform in single precision, and coordinates were saved periodically for analysis.

Trajectories were analyzed using MDAnalysis (30). Residue fluctuations were quantified as Cα root-mean-square fluctuations (RMSF). For each chain, frames were aligned to the chain-specific average structure, chain-specific RMSF values were calculated, and the three chains were averaged to obtain a per-residue mean RMSF profile. Backbone dihedral sampling was characterized by Ramachandran analysis with ϕ and ψ angles extracted for all nonterminal residues. Residue-specific Ramachandran plots were generated for selected sites of interest.

Interchain hydrogen-bond analysis was restricted to the classical collagen hydrogen bonds from the Gly amide NH of one chain to the carbonyl oxygen of the X-position residue on the adjacent chain. For each saved frame, candidate hydrogen bonds were identified using an N···O distance cutoff of 3.5 Å and a donor-hydrogen-acceptor angle cutoff of 125°. Only interchain contacts satisfying this canonical donor-acceptor pattern were retained. The total number of classical interchain hydrogen bonds was then counted for each frame of each trajectory. For each trajectory, framewise hydrogen-bond counts were averaged over the saved production frames and used for group summaries and statistical comparisons. Statistical comparisons were performed on peptide-level means using two-tailed Mann-Whitney U tests.

## Results

### 1. Bioinformatic analysis identifies a strong GP-associated sequence signal at pathogenic *COL3A1* glycine substitution sites

To determine whether pathogenic glycine substitutions in collagen occur preferentially in specific local sequence environments, we performed a sequence-context enrichment analysis of clinically reported mutations. For this analysis, we focused on type III collagen because of its homotrimeric structure and the extensive clinical variant data available for the associated disorder vascular Ehlers-Danlos syndrome (vEDS). Each reported case of a Gly missense mutation contributed one count to the analysis, so recurrent substitutions at the same glycine site were weighted proportionally. For each mutated glycine, we examined the eight flanking positions spanning two triplets upstream and two triplets downstream in the reference α1(III) chain sequence: *GX_-2_Y_-2_GX_-1_Y_-1_**G**X_1_Y_1_GX_2_Y_2_G*. Then we compared the observed case-weighted residue frequencies at each position with the corresponding background frequencies *f_i_(a)* across all repeating glycine residues within the main collagen domain of type III collagen sequence.

The distribution of flanking residues around pathogenic Gly substitution sites was clearly nonrandom (Figure 2). Sequence-context enrichment analysis identified 14 significantly enriched residue-position cells (q < 0.05 and fold-enrichment > 1). The strongest signal was Pro at X1, where Pro immediately follows the substituted Gly residue. This feature showed a case-weighted count of n = 301 and an FDR-adjusted q = 2.4 × 10⁻^11^ (Figure 2). Additional enriched positions included Thr at X_2_, Ser at Y_2_, and Asn and Pro at X_-1_, along with several lower-count enrichments at other positions (Figure 2). Thus, although pathogenic Gly substitutions in *COL3A1* were associated with multiple local sequence features, Pro immediately following the substituted Gly residue (GP context) emerged as the clearest and most heavily supported signal in the dataset. A similar analysis of Gly substitutions in the type II collagen gene *COL2A1* showed the same tendency toward increased Pro frequency at X1, although this enrichment did not reach statistical significance after FDR correction, likely reflecting the smaller available clinical dataset. The complete residue-position matrix, including observed case-weighted count *n_i_(a)*, the background frequency *f_i_(a)*, the fold-enrichment *[n_i_(a)/ N_i_]/ f_i_(a)*, and FDR-corrected *q*-values, is provided in Table S2.

**Figure 2.**
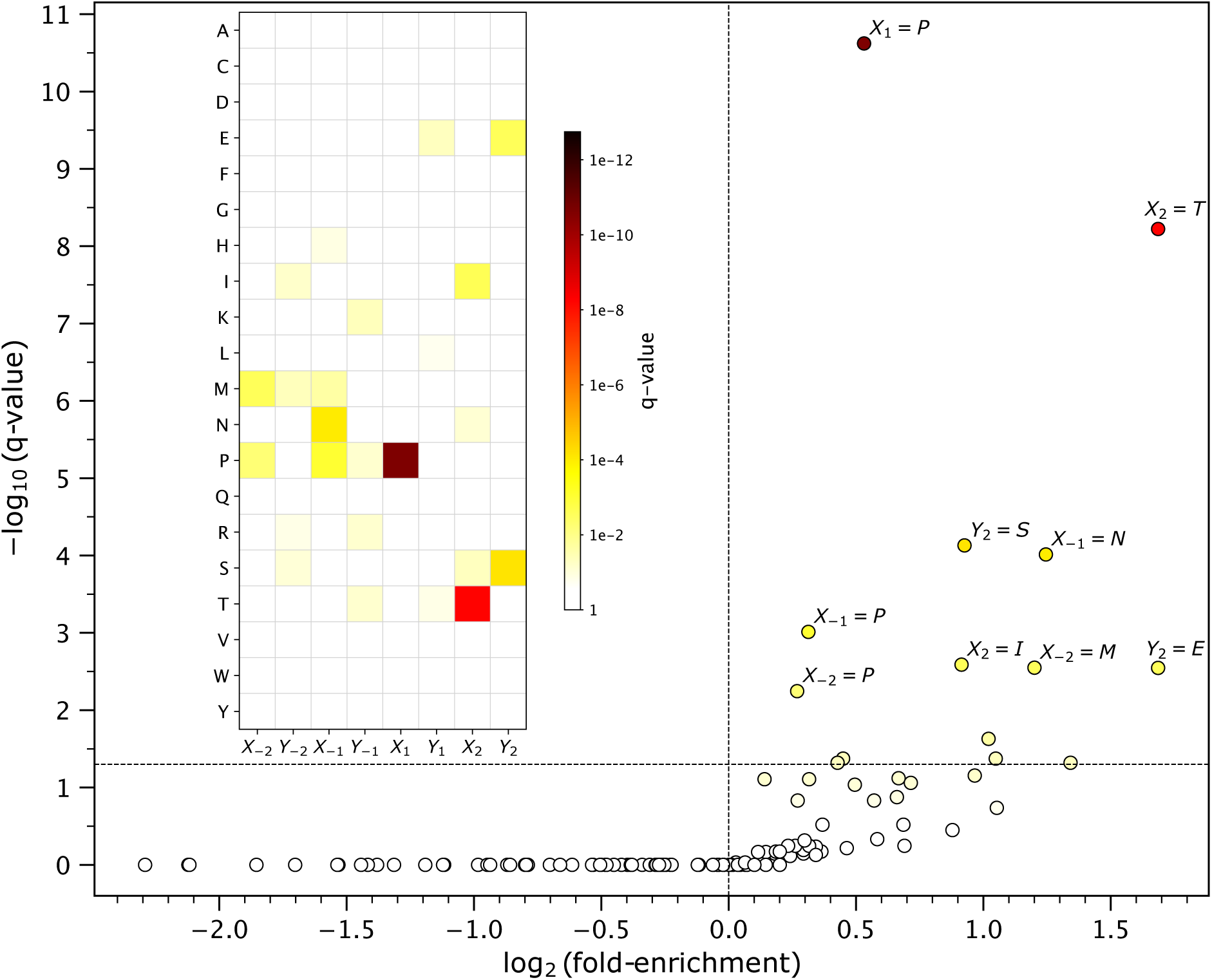
Case-weighted sequence-context enrichment analysis of pathogenic *COL3A1* glycine substitutions identifies Pro at X1 as the dominant local sequence signal. Volcano plot summarizing all 160 residue-position tests: each dot on the plot is a single amino acid in one of the eight flanking positions. The x-axis shows log_2_(fold-enrichment) relative to the corresponding background frequency in the reference α1(III) chain triple-helical sequence, and the y-axis shows −log_10_(*q*-value). Dashed lines indicate the enrichment threshold (fold-enrichment > 1) and the significance threshold (*q* < 0.05). Significantly enriched residue-position combinations are colored and labeled. Inset is a heatmap showing the FDR-adjusted *q*-value for each amino acid at eight flanking positions surrounding pathogenic glycine missense substitutions in the uninterrupted triple-helical region of α1(III) chain. Cells meeting the enrichment criterion after Benjamini-Hochberg correction (*q* < 0.05) are highlighted, with warmer colors indicating stronger significance. Pro at *X_1_*, corresponding to a GP context, was the most strongly enriched residue-position combination.

Because X_1_ = Pro has a direct structural interpretation in the collagen triple helix, this result provided the strongest bioinformatic rationale for focusing subsequent experiments on GP-associated sequence environments. The enrichment of pathogenic substitutions in this context is consistent with the idea that some local collagen sequence environments are less permissive to Gly replacement than others. Although the bioinformatic analysis alone does not establish mechanism, it identifies GP as the most compelling sequence-level candidate for the biophysical and molecular modeling tests described below.

### 2. CMP design with stabilizing termini enables analysis of glycine substitutions in native collagen sequence contexts

CMPs were designed to include a 4-triplet (12-residue) sequence derived from native collagen (Figure 1), which was expected to be sufficient to examine sequence-context effects, as suggested by the bioinformatics analysis. Unlike classical host-guest CMPs, in which a small guest sequence or a limited number of residues is varied within a repetitive (GPO)_n_ stabilizing host, the present approach preserves an entire native four-triplet collagen sequence around the mutation site. It was intended as a stable peptide model for testing pathogenic Gly substitutions in their native local sequence contexts. As described in the Materials and Methods section, annealing during peptide preparation was essential for the highly stable WT peptides to ensure proper in-register chain alignment. Very rapid cooling can produce stable misaligned intermediates during triple-helix association; these misfolded triple helices must dissociate to monomers before correct folding can begin, whereas annealing facilitates proper folding by reducing the population of such off-pathway states (25,31). The high stability of the WT peptides allowed introduction of Gly substitutions without the need for stabilizing agents such as trimethylamine N-oxide, as previously suggested (16). In this design, the (GPO)_4_ stabilizing segments at both the N- and C-termini ensured complete peptide folding even for strongly destabilizing substitutions such as G→R. Each sequence began and ended with Gly, consistent with either the (Gly-Xaa-Yaa) or (Xaa-Yaa-Gly) repeating notation. An additional Tyr residue was included at the C-terminus to permit accurate concentration determination, which is essential for biophysical measurements. N- and C-terminal blocking groups reduced end repulsion and minimized intermolecular association at physiological pH (Figure 1).

Within the OI set, peptides were selected to sample a broad range of clinically relevant sequence environments while enabling direct comparisons. The A1-G382S/A1-G382R pair provided a same-site comparison of G→S and G→R substitutions associated with milder and more severe OI phenotypes, respectively. Additional α1(I) G→S peptides, including A1-G448S, A1-G832S, A1-G862S, and A1-G964S, expanded the range of sequence environments and clinical outcomes represented. To examine the contribution of chain-specific sequence context, we also synthesized A1-G661S and the corresponding α2(I) sequence context, A2-G661S, allowing comparison of the same G→S substitution in two distinct type I collagen environments associated with severe OI and osteoporosis, respectively. The vEDS set included adjacent observed sites within the same WT sequence, T3-G16S and T3-G19S; same-site Ser and Arg substitutions at T3-G415 within a sequence containing the GVMGFO DDR-binding motif (32); and additional disease-associated sites, T3-G70R, T3-G277R, and T3-G619R (Table 1).

**Table 1.**
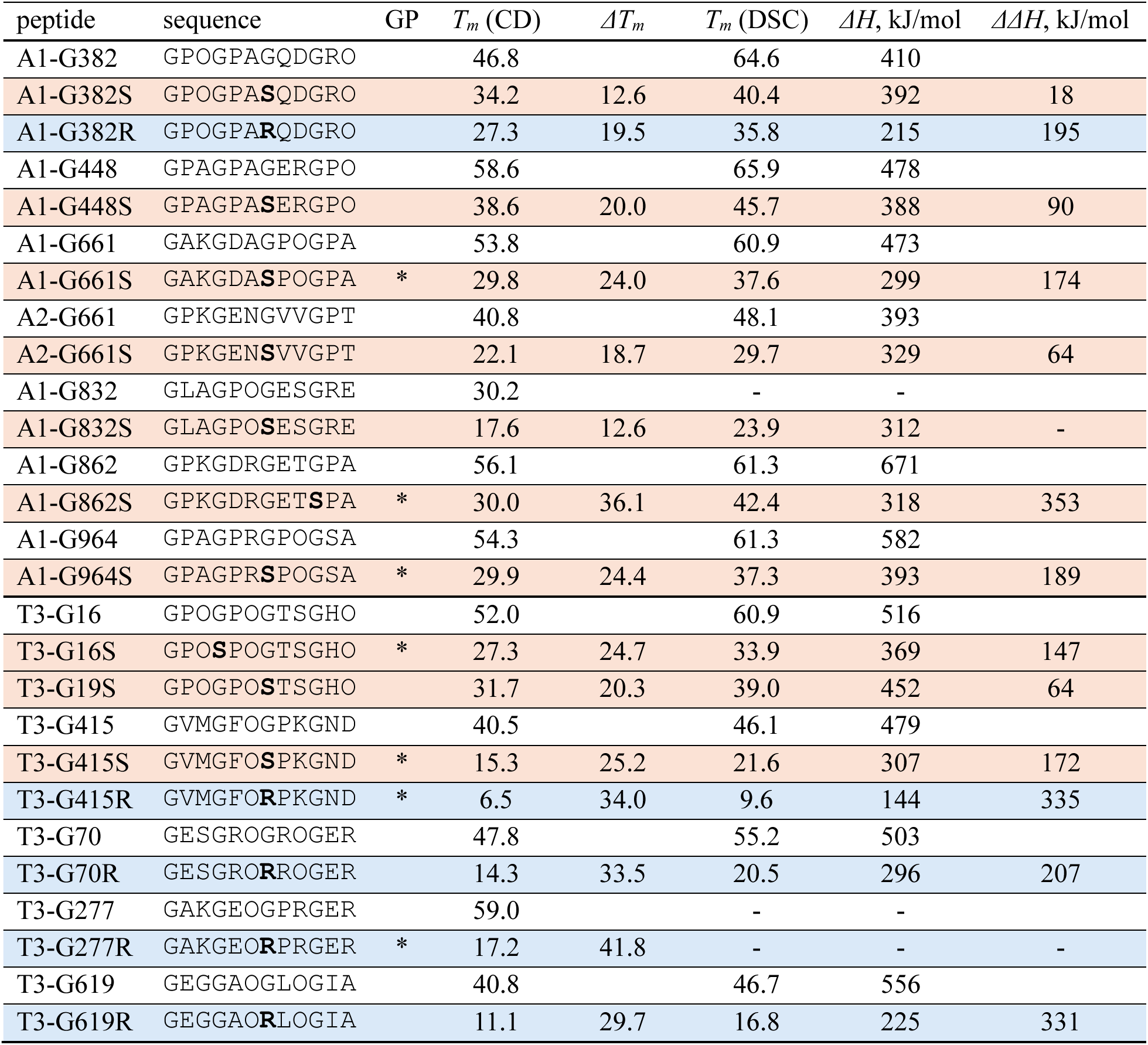
Collagen model peptides (CMPs) used in this study. All mutants (G→S with orange background and G→R with blue background) are matched with WT. The table summarizes peptide identity, middle sequence corresponding to yellow shading on Figure 1, mutant GP/nonGP classification, CD-derived *T_m_* at 0.1°C/min, DSC-derived *T_m_* at 1.0°C/min, destabilization *ΔTm*, calorimetric enthalpy for each WT and mutant peptide, and enthalpy loss *ΔΔH* for mutants. An asterisk indicates the GP context in mutant peptides. Dashes indicate samples for which DSC measurements were not available because sufficient purified peptide material was not available at the time of DSC analysis.

### 3. Biophysical measurements show that the consequences of glycine substitution depend on local sequence context

Biophysical characterization of the matched WT and mutant CMP pairs confirmed that the stabilizing-terminal design (Figure 1) enabled incorporation of 12-residue native collagen sequence windows containing G→S and G→R mutation sites while preserving an overall triple-helical conformation. As expected, DSC-derived transition temperatures were generally higher than CD-derived values, consistent with the faster scan rate, and CD-derived *T_m_* values were therefore used for cross-peptide comparisons of mutation-induced destabilization (Table 1). All peptides studied had *T_m_* values in the measurable range, spanning 6.5 °C for T3-G415R to 59 °C for T3-G277 (Table 1). For mutant peptides, *ΔT_m_ = T_m_(WT) – T_m_(mutant)* served as the primary measure of loss of thermal stability, whereas *ΔΔH* = *ΔH(WT) – ΔH(mutant)* from DSC provided a complementary measure of the enthalpic penalty associated with the substitution. Across the full peptide set, every glycine substitution reduced stability relative to its matched WT peptide, but the magnitude of both *ΔT_m_* and *ΔΔH* varied markedly among peptides, indicating strong effects of both the substituting residue and the local sequence environment (Table 1).

#### Gly→Arg substitutions caused larger destabilization than Gly→Ser substitutions

To assess the contribution of substitution identity independent of surrounding sequence, we compared G→S and G→R substitutions across the peptide set and at matched sites where both replacements were examined. Overall, G→R substitutions were more damaging than G→S substitutions. This difference is illustrated by the type III collagen peptide series centered at Gly415 (Figure 3A,B). Both vEDS-causing substitutions, T3-G415S and T3-G415R, were strongly destabilized relative to WT, but the G→R substitution caused the larger effect in both CD and DSC assays. The unfolding transition of T3-G415R was shifted to substantially lower temperature than that of T3-G415S, and DSC showed a correspondingly larger reduction in unfolding enthalpy (Table 1).

**Figure 3.**
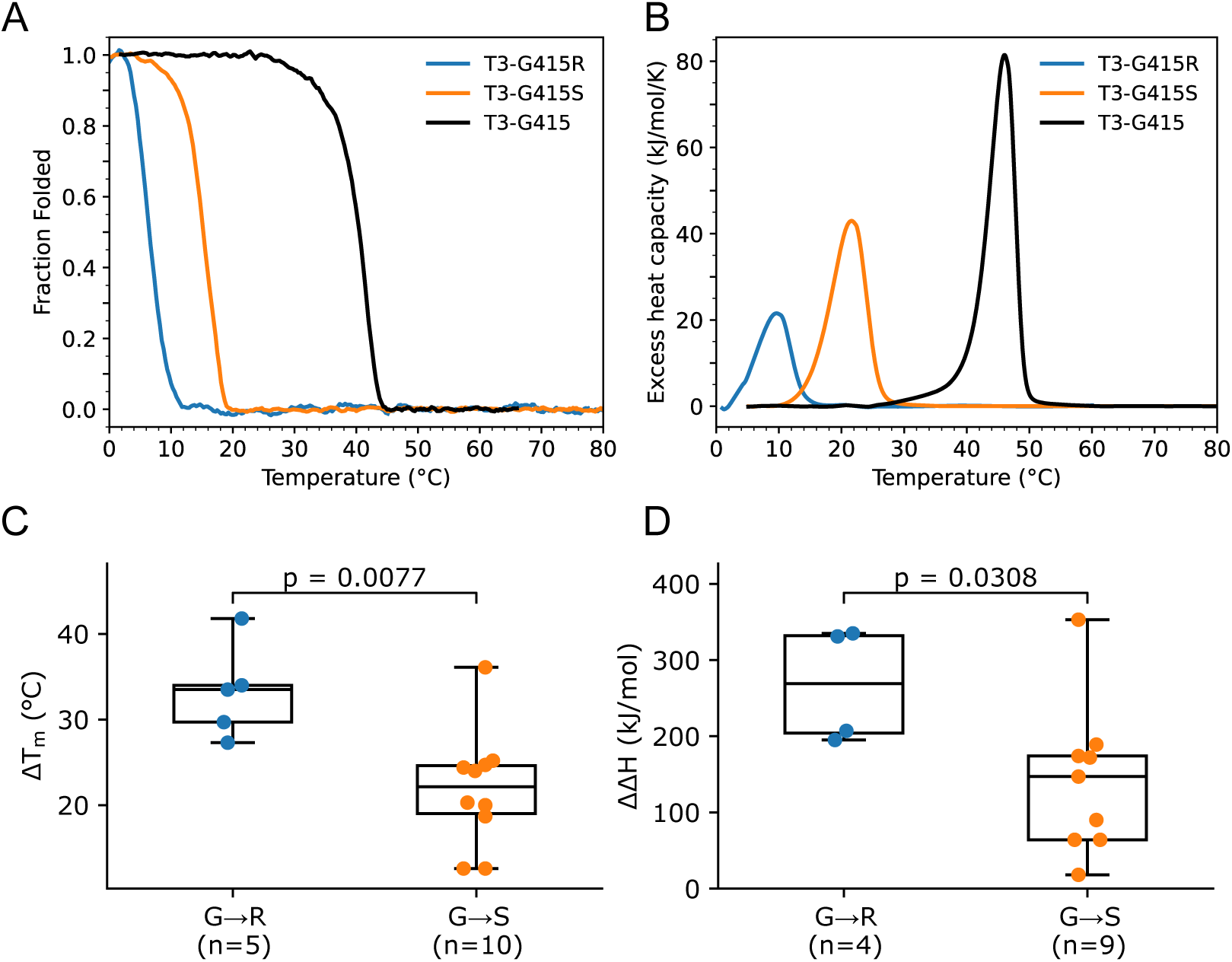
Gly→Arg substitutions are more destabilizing than Gly→Ser substitutions. Comparison of different replacing residues at the same collagen sequence position. (A) Representative CD thermal melts for T3-G415 WT, T3-G415S, and T3-G415R modeling the same mutation site in α1(III) chain. (B) DSC thermograms for the same peptide set. (C-D) Summary of *ΔT_m_* and *ΔΔH* for all same-site G→R vs. G→S comparisons of peptides included in the study. Replacement of Gly with Arg produced greater destabilization than replacement with Ser at the same local sequence position with two-tailed Mann-Whitney p-values shown on the plot.

The present CMP set reproduced that relationship across multiple native sequence contexts. The *ΔT_m_* values for G→S peptides ranged from 12.6 to 36.1 °C, whereas those for G→R peptides ranged from 27.3 to 41.8 °C. This difference was significant (p=0.0077) by a two-tailed Mann-Whitney test (Figure 3C). A similar relationship was observed for the enthalpic contribution to peptide stability. For peptides with measurable calorimetric transitions, G→R substitutions also showed larger *ΔΔH* values overall than G→S substitutions, with a significant (p=0.0308) difference by a two-tailed Mann-Whitney test (Figure 3D).

#### Gly→Ser substitutions at GP sites were more destabilizing than those at nonGP sites

The strongest sequence-context effect in the peptide set was observed when all Gly→Ser substitutions were grouped according to whether the mutated Gly was followed by Pro (GP context) or by another residue (nonGP context). This effect is illustrated directly by the paired type III collagen peptides T3-G16S and T3-G19S (Figure 4A,B), which provide a stringent same-context comparison because both mutant peptides had identical amino acid composition and were examined against the same WT peptide background. In this pair, Gly16 is followed by Pro (GP context), whereas Gly19 is followed by Thr, representing a nonGP site. In the native α1(III) sequence, this region is preceded by multiple GPO triplets, so the N-terminal (GPO)_4_ segment in this peptide closely matches the natural local sequence context rather than acting only as an artificial stabilizing host. Both substitutions destabilized the peptide relative to WT, but the GP-site mutation was more damaging: the CD and DSC melting transitions of T3-G16S occurred at lower temperatures than those of T3-G19S, and a larger loss of unfolding enthalpy was observed for T3-G16S, suggesting greater interchain hydrogen-bond disruption in the GP context.

**Figure 4.**
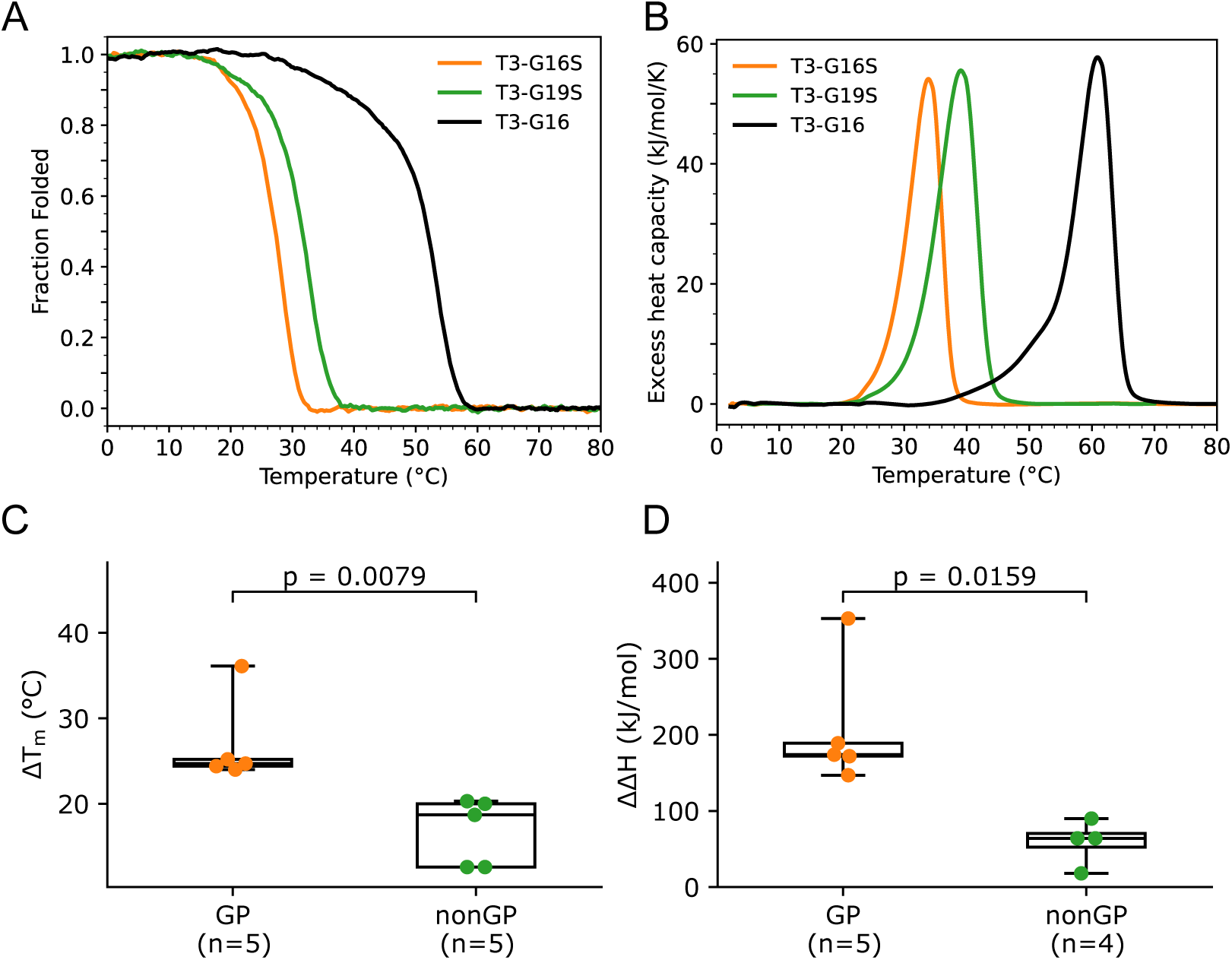
Gly→Ser substitutions are more destabilizing at GP sites than at nonGP sites. Representative biophysical comparisons showing that the effect of Gly→Ser substitution depends strongly on local sequence context. (A) CD thermal melts for the T3-G16 WT peptide and two mutants based on context: T3-G16S (GP) and T3-G19S (nonGP). (B) DSC thermograms for the same peptide set. (C-D) Summary of *ΔT_m_* and *ΔΔH* for G→S peptides in the GP context vs. G→S peptides in nonGP context. Replacement of Gly with Ser caused significantly greater destabilization when the mutated Gly was immediately followed by Pro with two-tailed Mann-Whitney p-values shown on the plot.

A direct comparison was also made between the type I collagen peptides A1-G661S and A2-G661S, which represent the same glycine position in the α1(I) and α2(I) chains, respectively. Although native type I collagen is a heterotrimer, the peptide system used here consists of homotrimeric CMPs built from either the α1(I) or α2(I) local sequence context. Within this simplified framework, the α1(I)-derived peptide was more strongly destabilized by the G→S substitution than the α2(I)-derived peptide. Specifically, A1-G661S showed a larger loss in thermal stability and unfolding enthalpy than A2-G661S, with *ΔT_m_* values of 24.0 and 18.7 °C and *ΔΔH* values of 174 and 64 kJ/mol, respectively. Notably, A1-G661S is a GP-site substitution.

The same ordering seen in this representative pair was reproduced across the broader peptide set, where the GP-site group was consistently shifted toward larger destabilization. For thermal stability, the median *ΔT_m_* was 24.7 °C for GP-site substitutions and 18.7 °C for nonGP substitutions, with a significant difference (p=0.0079) by a two-tailed Mann-Whitney test (Figure 4C). The same pattern was seen for calorimetric enthalpy, where GP-site substitutions also showed larger *ΔΔH* values overall than nonGP substitutions (Figure 4D). Because DSC measurements were unavailable for two WT peptides, the ΔΔH comparison was performed on the subset with measurable calorimetric transitions, but the separation between GP and nonGP groups remained clear and significant (p=0.0159).

Together, these measurements show that the outcome of glycine substitution in collagen depends on both the identity of the replacing residue and the immediate sequence environment around the mutation site. G→R substitutions were more destabilizing than G→S substitutions, but the strongest sequence-context effect in the present peptide set was the greater destabilization produced by G→S substitutions when the mutated Gly was followed by Pro.

### 4. Molecular modeling links restrictive sequence environments to greater hydrogen-bond loss and reduced backbone accommodation

To examine the structural basis for the sequence dependence observed experimentally, we performed all-atom molecular dynamics simulations on the same WT and mutant CMPs in explicit CHARMM TIP3P water at 300 K using OpenMM and the CHARMM36mGP force field (29). Across all WT and mutant CMPs analyzed, the overall triple-helical fold was retained over 500 ns trajectories, indicating that the mutation-dependent differences described below reflect local accommodation within the triple helix rather than global unfolding. Root mean square fluctuations (RMSF) were highest at the peptide termini and in the central region surrounding the mutation site and were lowest in the middle of the (GPO)_4_ subsequences, yielding a W-shaped RMSF profile along the peptide sequence (Figure S1).

We next asked how Gly substitutions in different sequence contexts perturb the interchain hydrogen-bonding network that stabilizes the collagen triple helix. For this analysis, only the canonical interchain hydrogen bonds from Gly-NH on one chain to the carbonyl oxygen of the X-position residue on an adjacent chain were counted, with 36 such bonds representing the ideal maximum for an intact 12-triplet triple helix. A local structural comparison of the A1-G661 WT and A1-G661S trajectories illustrates the physical basis for hydrogen-bond loss at a GP-site G→S mutation. In the WT peptide, the central region retained the regular ladder of canonical interchain backbone hydrogen bonds characteristic of the collagen triple helix, whereas in the mutant peptide several such bonds were lost near the substitution site (Figure 5A). We observed the same overall tendency across the full peptide set: Gly substitutions generally produced a localized loss of canonical interchain hydrogen bonds centered at the mutation site, with limited propagation to neighboring triplets in most cases (Figure S2).

**Figure 5.**
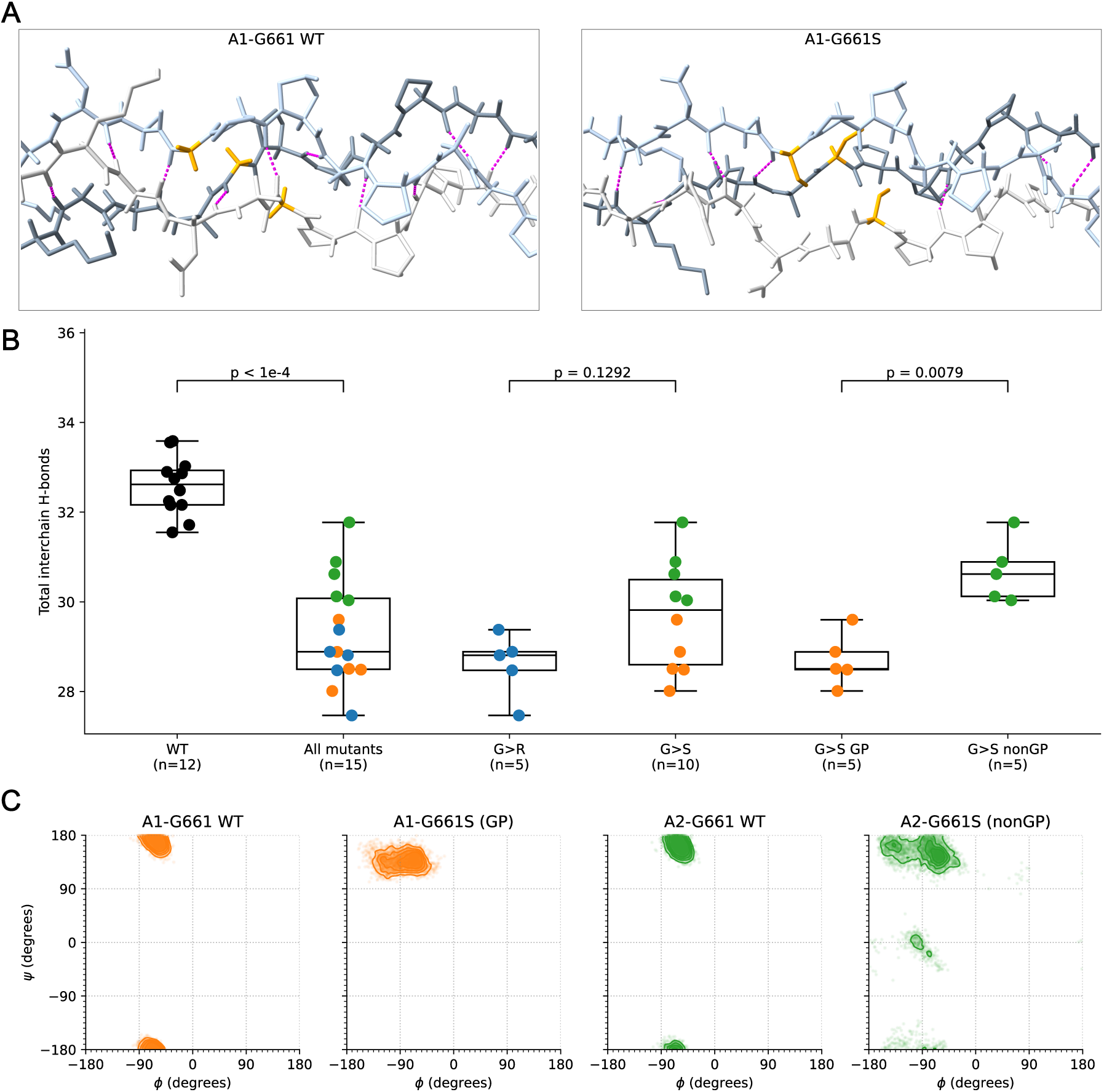
Molecular modeling links restrictive sequence environments to greater hydrogen-bond loss and reduced backbone accommodation. **(A)** Representative structural snapshots of A1-G661 WT (left) and A1-G661S (right) centered on residue 661. The three peptide chains are shown in distinct shades of gray, the residue at position 661 is highlighted in orange, and canonical interchain backbone hydrogen bonds are shown as magenta dashed lines. **(B)** Mean numbers of canonical interchain hydrogen bonds from 500 ns molecular dynamics trajectories for WT peptides, all mutants, G→R mutants, all G→S mutants, GP-site G→S mutants, and nonGP G→S mutants. Box plots show peptide-level distributions; points represent individual peptides and are colored by class (WT, black; G→R, blue; GP-site G→S, orange; nonGP G→S, green). *P*-values were calculated using two-tailed Mann-Whitney U tests. **(C)** Residue-specific Ramachandran plots for A1-G661/A1-G661S (GP context) and A2-G661/A2-G661S (nonGP context), showing pooled backbone dihedral angles from all three chains over the production trajectories. The nonGP mutant samples broader conformational space than the GP-site mutant.

Across the full peptide set, WT peptides clustered near the expected maximum, whereas mutant peptides showed reduced hydrogen-bond counts overall (Figure 5B and Table S3). When all mutants were considered together, peptide-level mean counts of classical interchain hydrogen bonds were significantly lower than in WT peptides (two-tailed Mann-Whitney U test, p < 0.0001), indicating that glycine substitution broadly weakens the canonical collagen hydrogen-bonding network. Stratification by mutation class and local sequence context revealed the same hierarchy observed experimentally. G→R mutants had lower hydrogen-bond counts than G→S mutants overall, consistent with greater structural disruption by arginine substitution, although this difference did not reach significance in the present peptide set (two-tailed Mann-Whitney U test, p = 0.1292). In contrast, the sequence-context effect within the G→S class was pronounced: GP-site G→S mutants showed significantly fewer canonical interchain hydrogen bonds than nonGP G→S mutants (two-tailed Mann-Whitney U test, p = 0.0079). Thus, even in trajectories that remained globally triple-helical, local sequence environment strongly influenced how Gly substitutions disrupt the canonical interchain hydrogen-bonding network.

Ramachandran analysis further clarified how local backbone accommodation of glycine substitution depends on sequence context (Figure 5C). In the WT A1-G661 and A2-G661 peptides, the residue-specific φ/ψ distributions at this site were similar despite the different local sequence environments, indicating that both peptides retained the characteristic triple-helical geometry in the absence of mutation. After G→S substitution, however, the two contexts diverged markedly. The GP-site mutant A1-G661S remained confined to a relatively restricted region of Ramachandran space, whereas the nonGP mutant A2-G661S sampled a substantially broader φ/ψ distribution, including additional low-occupancy conformational states. Residue-specific Ramachandran plots across all simulated systems showed the same general pattern of mutation-dependent shifts in local backbone sampling, consistent with the examples highlighted in Figure 5C (Figure S3). Thus, the nonGP environment appears to be better able to adjust local backbone geometry to accommodate the serine substitution, whereas the more rigid GP environment remained more conformationally constrained, resulting in greater distortion of the stabilizing hydrogen-bond network. Together, the MD simulations provide a structural explanation for the sequence dependence observed in both the bioinformatic and peptide analyses: mutations in restrictive local environments, especially GP sites, show greater loss of classical interchain hydrogen bonds and poorer local backbone accommodation than mutations in more permissive contexts.

## Discussion

A central question in collagen mutation biology is why glycine substitutions, all of which violate the same structural requirement of the triple helix, differ so markedly in their phenotypic consequences. Earlier studies established that the structural effects of missense mutations are influenced both by the identity of the replacing residue and by the local sequence flanking the Gly site (16,33). The present results confirm these relationships in newly designed CMPs, but the clearest new finding is that local sequence context, particularly GP versus nonGP context, strongly influences how a Gly substitution is accommodated within the triple helix. In the peptide set examined here, GP context emerged as the strongest and most reproducible determinant of enhanced destabilization for G→S substitutions, and the same pattern appeared independently in the case-weighted sequence analysis, peptide thermodynamic measurements, and molecular dynamics simulations.

A major strength of the present study is the peptide design itself. Modeling pathogenic Gly substitutions in short CMPs is challenging because such substitutions can strongly disrupt folding, and earlier mutation-bearing peptides often remained incompletely folded or required stabilizing agents to form a measurable triple helix (16,33). Here, a native four-triplet collagen sequence was placed between two stabilizing (GPO)_4_ segments, allowing the mutation site to be examined within a well-folded triple-helical scaffold (Figure 1). Terminal blocking groups reduced end effects and higher-order association, and the annealing protocol helped avoid misregistered assemblies in the most stable WT peptides. The cooperative CD and DSC transitions indicate that this design provides a robust platform for comparing local sequence effects on glycine substitution while preserving a native collagen-derived sequence environment.

The resulting peptide set is also useful because it functions as a comparative panel rather than simply a collection of individual mutants. It spans multiple regions of type I and type III collagen and enables several controlled comparisons within a single experimental framework, including G→S versus G→R substitutions at the same site, the same nominal G→S substitution in different chain contexts, and neighboring Gly sites within the same WT sequence. In that sense, these CMPs provide experimentally controllable models for clinically observed collagen mutations while preserving the native local sequence within a defined triple-helical scaffold. At the same time, they remain intentionally simplified systems and do not reproduce the heterozygous mutation state, the heterotrimeric architecture of type I collagen, C-propeptide-directed assembly, or longer-range structural and matrix-level effects (34,35). Therefore, the present CMPs are best interpreted as controlled models for isolating local sequence-context effects on Gly substitution tolerance, rather than as full physiological models of dominant collagen mutations.

Within the α1(I) OI subset, a qualitative trend was also evident: mutations associated with more severe reported OI phenotypes tended to produce greater destabilization, although the current sample size is too limited for formal statistical analysis. Specifically, mutations reported in severe OI types II and III, including A1-G382R, A1-G661S, A1-G862S, and A1-G964S, produced larger decreases in peptide stability (*ΔT_m_* = 19.5-36.1 °C) than mutations reported in milder type IV OI cases, including A1-G382S, A1-G448S, and A1-G832S (*ΔT_m_* = 12.6-20.0 °C) (Table 1). This is consistent with the idea that more severe OI mutations tend to impose greater triple-helix destabilization.

The present study further shows that the greater destabilizing effect of G→R relative to G→S, previously observed mainly in repetitive host-guest peptides under stabilizing conditions, also holds in native collagen sequence contexts. Across the sequence environments examined, G→R substitutions generally caused greater destabilization than G→S substitutions, in agreement with earlier peptide studies (16). The clearest evidence comes from direct same-site comparisons in the A1-G382S/R and T3-G415S/R pairs, where CD and DSC showed that Arg caused a further 7-9 °C decrease in *T_m_* and a 160-180 kJ/mol greater loss of unfolding enthalpy relative to Ser (Figure 3A and Table 1). Thus, the present CMP system not only confirms the established residue hierarchy, but does so in natural collagen-derived sequence environments and across multiple mutation sites. The hydrogen-bond analysis did not show a statistically significant separation between the G→R and G→S groups, likely because of limited sample size (Figure 5B), but the thermal measurements clearly distinguished the two classes (Figure 3C,D). At the same time, the distributions overlapped, indicating that a G→S mutation in a restrictive local environment can be as damaging as, or even more damaging than, a G→R mutation in a more permissive one. A hierarchy of replacing residues remains useful and necessary, but it is not sufficient.

The bioinformatic analysis provided an important clue to the nature of that restrictive environment. Among pathogenic *COL3A1* glycine substitutions, Pro at the *X_1_* position, corresponding to a GP context, emerged as the dominant enriched local sequence feature (Figure 2). This result is striking because Pro-rich sequences are generally associated with favorable WT triple-helix formation (23,26,36). The present findings therefore indicate that WT stability and mutational tolerance are not equivalent properties. A local sequence can stabilize the native triple helix but still tolerate loss of the obligatory Gly residue poorly. In that sense, enrichment of GP-associated sites in *COL3A1* appears to mark not a weak region of the WT helix, but rather a region of reduced tolerance to mutation.

This conclusion is directly supported by the matched A1-G661S and A2-G661S comparison. These peptides model the same nominal glycine position in the α1(I) and α2(I) chains, but the local sequence environments are different. When the G→S substitution was introduced, both peptides were destabilized relative to WT, but A1-G661S showed a larger loss of thermal stability and calorimetric enthalpy than A2-G661S (Table 1). In the simulations, the GP-site mutant A1-G661S also showed a larger reduction in classical interchain hydrogen bonding than the nonGP mutant A2-G661S. Ramachandran analysis further showed that after substitution the nonGP context sampled a broader φ/ψ distribution, whereas the GP context remained more restricted (Figure 5C). Thus, within this homotrimeric CMP system, the α1(I)-derived context was less permissive than the α2(I)-derived context. This difference is consistent with the local sequence itself: in the α1(I) context, the mutated Gly is followed by Pro, whereas in the α2(I) context it is followed by Val. The clinical contrast between severe OI reported for α1(I) G661S and osteoporosis reported for α2(I) G661S is directionally consistent with these biophysical differences, although this comparison should be interpreted cautiously because native type I collagen contains two α1(I) chains and only one α2(I) chain.

An even cleaner demonstration of local sequence effects is provided by the T3-G16S and T3-G19S pair. These two mutant peptides have identical amino acid composition and were examined against the same WT background, so they differ mainly in the position of the mutation within the same sequence. The key difference between these two peptides was that Gly16 is followed by Pro, whereas Gly19 is followed by Thr. T3-G16S was more destabilizing than T3-G19S, with a 3.4 °C lower *T_m_* and an approximately 80 kJ/mol greater loss of unfolding enthalpy (Figure 4A; Table 1). Because these peptides do not differ in replacing residue or in overall composition, the greater destabilization of T3-G16S cannot be attributed to a more damaging side chain or to a different peptide background. Instead, it points to a genuine positional effect within the local collagen sequence. The broader comparison across all G→S peptides supports the same interpretation: as a group, GP-site substitutions produced larger losses of thermal stability and unfolding enthalpy than nonGP substitutions (Figure 4C,D).

The molecular dynamics simulations provide a structural explanation for this behavior. Across all WT and mutant CMPs, the overall triple-helical fold was retained throughout the 500 ns trajectories, indicating that the experimentally observed differences do not require global unfolding of the peptide. Instead, the main effects were local. GP-site G→S substitutions retained fewer canonical interchain hydrogen bonds than nonGP substitutions, and the matched A1-G661S/A2-G661S comparison showed that the nonGP context could sample a broader range of local backbone conformations after mutation while preserving a larger fraction of the canonical hydrogen-bond network. In this case, broader conformational sampling does not mean a more native local structure, but rather greater local accommodation of the unfavorable Ser substitution. By contrast, the GP context appears less able to adjust local backbone geometry after loss of Gly, resulting in greater disruption of the stabilizing hydrogen-bond network. These results support the view that Pro-adjacent sites are relatively intolerant because they are less adaptable in the mutant state.

This interpretation helps reconcile two features of collagen sequence that might otherwise seem contradictory. Pro and Hyp are classical stabilizing residues in collagen, and triple-helix propensity studies, stability algorithms, and recent CMP work all underscore the strong sequence dependence of WT collagen stability (26,36,37). In the present peptide set, several mutations occur near regions associated with triple-helix stabilization, including GPO-rich segments and KGE/KGD-containing charged motifs. Recent work also suggests that interchain salt bridges can remain intact in the vicinity of triple-helix perturbations such as Gly substitutions (38). At the same time, the present results indicate that a Pro immediately following the mutated Gly can make that site less tolerant of substitution. Thus, the sequence features that stabilize the WT helix are not necessarily the features that best tolerate perturbation. Distinguishing WT stability from mutational tolerance may therefore be important for predicting mutation outcome.

The present findings also have implications for interpretation of collagen disease mutations. For OI, the importance of chain identity, substituting residue, and position along the molecule has long been recognized, but prediction of outcome from sequence remains incomplete (2,33,39). For vEDS, genotype-phenotype relationships are also complex and are influenced by variant class and molecular mechanism in addition to local sequence features (4,5,40,41). The current results suggest that immediate sequence context should be considered as an additional mechanistic variable in both settings. At the same time, local sequence alone is unlikely to determine clinical outcome, because intracellular proteostasis and procollagen handling can also contribute substantially (9,42,43). Nevertheless, the agreement among the bioinformatic, biophysical, and modeling analyses strengthens the conclusion that GP context is not an isolated feature of one assay, but a robust determinant of mutational intolerance.

In summary, the present data support a simple but important conclusion: the consequence of a collagen glycine substitution depends not only on the identity of the replacing residue, but also on the permissiveness of the surrounding sequence environment. The immediate sequence environment surrounding a glycine site behaves as a local structural unit that influences how strongly the mutation perturbs hydrogen bonding, backbone geometry, and thermal stability. In the current dataset, GP context emerged as the clearest example of a restrictive local environment. The main advance of this study is the demonstration that a following Pro can strongly modulate the outcome of glycine substitution, such that selected G→S mutations in GP contexts can approach the destabilizing effect of G→R substitutions. Together, the bioinformatic, biophysical, and modeling results support a framework in which GP versus nonGP context is a major determinant of collagen mutational outcome.

## Data availability

All data supporting the findings of this study are included in the article, its Supplemental Material, and the associated Zenodo datasets. Complete RP-HPLC chromatograms and MALDI-TOF mass spectra for all synthesized peptides are available in Zenodo at DOI: 10.5281/zenodo.20560423. Molecular dynamics input files for all simulated CMPs are available in Zenodo at DOI: 10.5281/zenodo.20750355.

## Acknowledgments

The author thanks Barbara Brodsky for more than 25 years of inspiration in collagen triple-helix research, for many stimulating discussions, and for access to circular dichroism and differential scanning calorimetry instrumentation. Support for preparation of osteogenesis imperfecta and vascular Ehlers-Danlos syndrome collagen model peptides was provided by the Osteogenesis Imperfecta Foundation and the National Organization for Rare Disorders. Computations reported in this work were performed using resources made available by the Flatiron Institute, a division of the Simons Foundation.

## Author contributions

A.V.P. designed the research, performed the research, analyzed the data, and wrote the manuscript.

