## Supplemental Figures and Tables for "Local sequence context determines the effect of glycine substitutions in collagen triple helices"

**Figure S1. Per-residue RMSF profiles for all WT and mutant peptides from 500 ns molecular dynamics trajectories.**
RMSF values are plotted against peptide residue number for each of the three chains separately, with Chain A shown in blue, Chain B in orange, and Chain C in green; the chain-averaged RMSF is shown as a thicker red line. Panels are arranged by collagen type and mutation status: type I WT, type I mutants, type III WT, and type III mutants. Corresponding WT and mutant peptides are positioned for direct comparison.

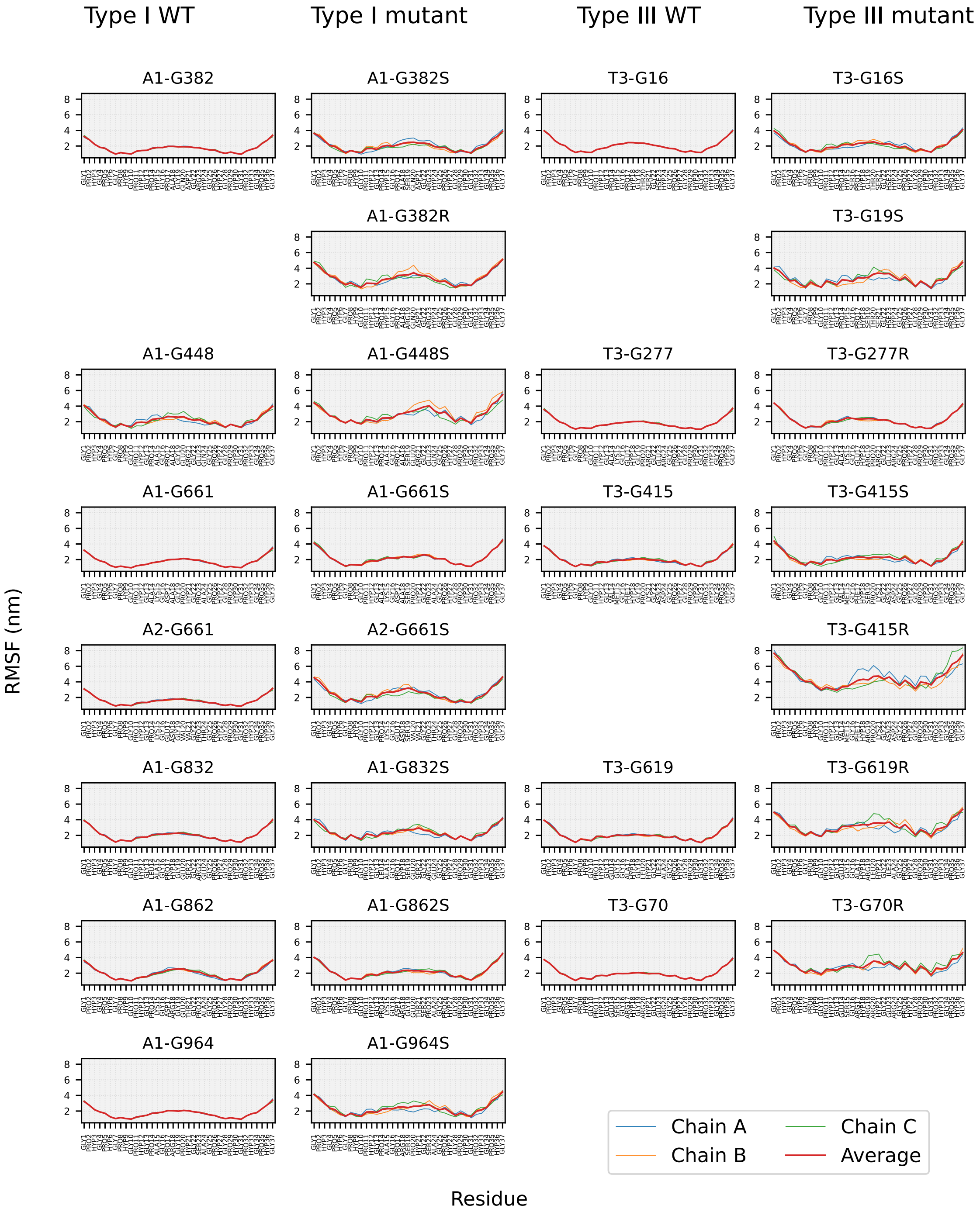


Figure S1. Per-residue RMSF profiles for all WT and mutant peptides

**Figure S2. Residue-by-residue classical interchain hydrogen-bond occupancy for all WT and mutant CMPs from 500 ns MD trajectories.**

Classical interchain hydrogen-bond occupancy is plotted as a function of Gly position along each peptide, allowing direct visualization of local loss of canonical Gly–Xaa hydrogen bonds after Gly substitution. For each Gly position, occupancy was calculated from the canonical interchain Gly–NH to X-position carbonyl O hydrogen bonds and averaged across the three chains over the trajectory. Panels are arranged by collagen type and mutation status: type I WT, type I mutants, type III WT, and type III mutants. Corresponding WT and mutant peptides are positioned for direct comparison.

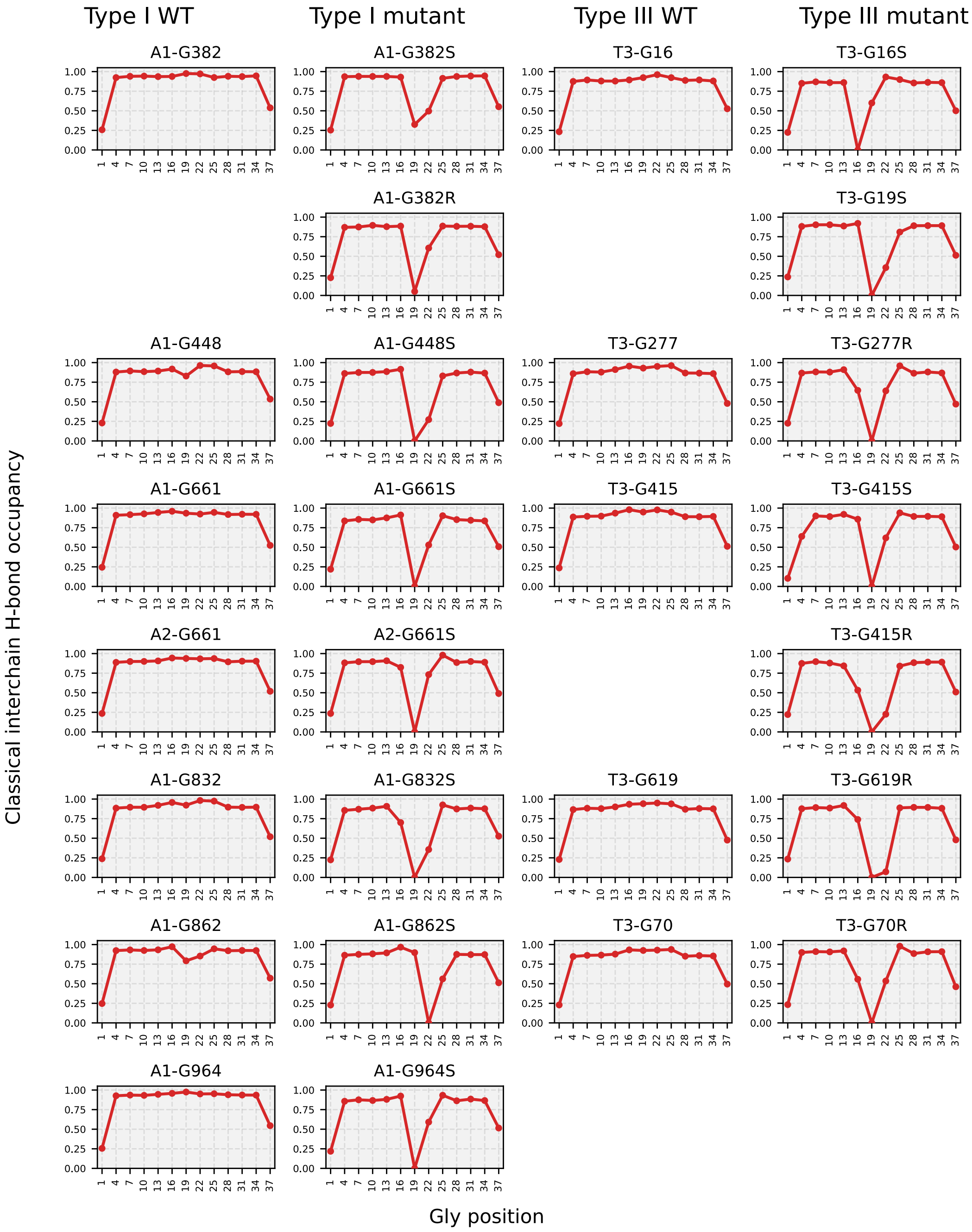


Figure S2. Residue-by-residue classical interchain hydrogen-bond occupancy

**Figure S3. Residue-specific Ramachandran plots for the mutation-site residue in all simulated systems.** Ramachandran distributions are shown for the residue at the mutation site in each WT and mutant peptide, extending the comparison presented in Figure 5C to the full peptide set. For each panel, ϕ/ψ values from all three chains were pooled over the 500 ns trajectory and plotted as residue-specific conformational distributions. WT peptides are shown in grey, Gly→Ser mutants in orange, and Gly→Arg mutants in blue. Panels are arranged by collagen type and mutation status: type I WT, type I mutants, type III WT, and type III mutants, with corresponding WT and mutant peptides positioned for direct comparison.

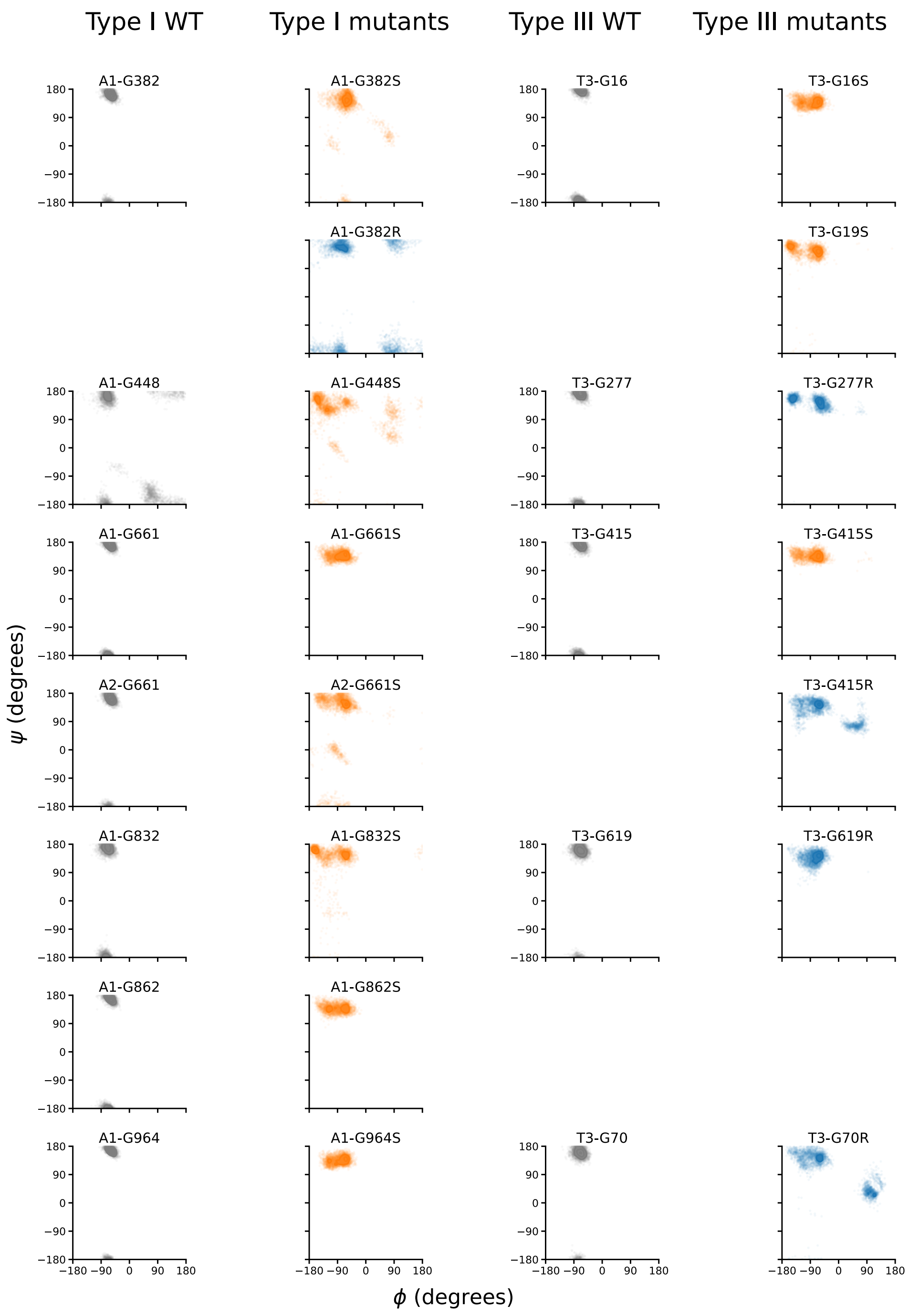


Figure S3. Residue-specific Ramachandran plots for the mutation-site residue

**Table S1. Peptide identity and purity data for all synthesized collagen model peptides.**

All peptides were synthesized by solid-phase Fmoc chemistry and purified to >95% by reverse-phase HPLC prior to biophysical analysis. Peptide identity was confirmed by MALDI-TOF mass spectrometry with external calibration. Calculated m/z values are reported for the assigned [M+H]$^+$ ion. Observed m/z values represent the [M+H]$^+$ ion detected by MALDI-TOF. Mass deviation (Δm, ppm) is reported as |calc − obs| / calc × $10^6$. Deviations of 3–217 ppm across the 27-peptide set are consistent with the expected performance of MALDI-TOF under external calibration conditions for peptides in the 3300–3800 Da range (typically in the tens to low hundreds of ppm). Complete raw characterization data (RP-HPLC chromatograms and MALDI-TOF spectra for all peptides) are deposited at Zenodo: DOI 10.5281/zenodo.20560423.

| peptide | sequence | calculated [M+H]$^+$ mono. (m/z) | observed MALDI-TOF (m/z) | ion | Δm (ppm) | purity % |
|---|---|---|---|---|---|---|
| A1-G382 | GPOGPOGPOGPOGPOGPAGQDGROGPOGPOGPOGPOGY | 3537.76 | 3537.59 | [M+H]$^+$ | 48 | 96.1 |
| A1-G382S | GPOGPOGPOGPOGPOGPA**S**QDGROGPOGPOGPOGPOGY | 3567.79 | 3567.65 | [M+H]$^+$ | 39 | 97.6 |
| A1-G382R | GPOGPOGPOGPOGPOGPA**R**QDGROGPOGPOGPOGPOGY | 3636.90 | 3636.51 | [M+H]$^+$ | 107 | 98.6 |
| A1-G448 | GPOGPOGPOGPOGPAGPAGERGPOGPOGPOGPOGY | 3478.74 | 3478.61 | [M+H]$^+$ | 37 | 96.3 |
| A1-G448S | GPOGPOGPOGPOGPAGPA**S**ERGPOGPOGPOGPOGY | 3508.77 | 3508.01 | [M+H]$^+$ | 217 | 99.9 |
| A1-G661 | GPOGPOGPOGPOGAKGDAGPOGPAGPOGPOGPOGY | 3410.66 | 3410.69 | [M+H]$^+$ | 9 | 97.1 |
| A1-G661S | GPOGPOGPOGPOGAKGDA**S**POGPAGPOGPOGPOGY | 3440.69 | 3440.65 | [M+H]$^+$ | 12 | 99.9 |
| A2-G661 | GPOGPOGPOGPOGPKGENGVVGPTGPOGPOGPOGY | 3511.81 | 3511.80 | [M+H]$^+$ | 3 | 99.9 |
| A2-G661S | GPOGPOGPOGPOGPKGEN**S**VVGPTGPOGPOGPOGY | 3541.84 | 3541.87 | [M+H]$^+$ | 9 | 99.8 |
| A1-G832 | GPOGPOGPOGPOGLAGPOGESGREGPOGPOGPOGY | 3542.78 | 3542.89 | [M+H]$^+$ | 31 | 95.7 |
| A1-G832S | GPOGPOGPOGPOGLAGPO**S**ESGREGPOGPOGPOGY | 3572.81 | 3572.67 | [M+H]$^+$ | 39 | 99.9 |
| A1-G862 | GPOGPOGPOGPOGPKGDRGETGPAGPOGPOGPOGY | 3321.57 | 3321.27 | [M+H]$^+$ | 90 | 99.9 |
| A1-G862S | GPOGPOGPOGPOGPKGDRGET**S**PAGPOGPOGPOGY | 3351.59 | 3351.23 | [M+H]$^+$ | 107 | 97.9 |
| A1-G964 | GPOGPOGPOGPOGPAGPRGPOGSAGPOGPOGPOGY | 3436.70 | 3436.95 | [M+H]$^+$ | 73 | 95.4 |
| A1-G964S | GPOGPOGPOGPOGPAGPR**S**POGSAGPOGPOGPOGY | 3466.73 | 3466.05 | [M+H]$^+$ | 196 | 99.8 |
| T3-G16 | GPOGPOGPOGPOGPOGPOGTSGHOGPOGPOGPOGY | 3505.72 | 3505.58 | [M+H]$^+$ | 40 | 99.9 |
| T3-G16S | GPOGPOGPOGPOGPO**S**POGTSGHOGPOGPOGPOGY | 3535.75 | 3535.32 | [M+H]$^+$ | 122 | 99.9 |
| T3-G19S | GPOGPOGPOGPOGPOGPO**S**TSGHOGPOGPOGPOGY | 3535.75 | 3535.33 | [M+H]$^+$ | 119 | 99.9 |
| T3-G415 | GPOGPOGPOGPOGVMGFOGPKGNDGPOGPOGPOGY | 3591.92 | 3591.84 | [M+H]$^+$ | 22 | 99.9 |
| T3-G415S | GPOGPOGPOGPOGVMGFO**S**PKGNDGPOGPOGPOGY | 3621.94 | 3621.36 | [M+H]$^+$ | 160 | 97.3 |
| T3-G415R | GPOGPOGPOGPOGVMGFO**R**PKGNDGPOGPOGPOGY | 3691.05 | 3691.72 | [M+H]$^+$ | 182 | 95.9 |
| T3-G70 | GPOGPOGPOGPOGESGROGROGERGPOGPOGPOGY | 3686.92 | 3686.54 | [M+H]$^+$ | 103 | 98.4 |
| T3-G70R | GPOGPOGPOGPOGESGRO**R**ROGERGPOGPOGPOGY | 3786.05 | 3786.55 | [M+H]$^+$ | 132 | 99.5 |
| T3-G277 | GPOGPOGPOGPOGAKGEOGPRGERGPOGPOGPOGY | 3626.91 | 3626.73 | [M+H]$^+$ | 50 | 97.7 |
| T3-G277R | GPOGPOGPOGPOGAKGEO**R**PRGERGPOGPOGPOGY | 3726.04 | 3726.55 | [M+H]$^+$ | 137 | 98.0 |
| T3-G619 | GPOGPOGPOGPOGEGGAOGLOGIAGPOGPOGPOGY | 3427.69 | 3427.18 | [M+H]$^+$ | 149 | 98.9 |
| T3-G619R | GPOGPOGPOGPOGEGGAO**R**LOGIAGPOGPOGPOGY | 3526.82 | 3526.84 | [M+H]$^+$ | 6 | 99.4 |

**Table S2. Complete residue-position matrix for case-weighted sequence-context enrichment analysis of pathogenic *COL3A1* glycine substitutions.**

Each row corresponds to one amino acid at one flanking position relative to the substituted Gly residue ($X_{-2}$, $Y_{-2}$, $X_{-1}$, $Y_{-1}$, $X_1$, $Y_1$, $X_2$, or $Y_2$). "observed_count" is the observed case-weighted count $n_i(a)$ for amino acid $a$ at position $i$. "background_frequency" is the fraction $f_i(a)$ of eligible COL3A1 reference Gly sites with residue $a$ at position $i$. "expected_count" is $E_i(a) = N_i f_i(a)$. fold_enrichment is $n_i(a)/E_i(a)$. "p_value" is the one-sided upper-tail binomial $p$-value for enrichment, and "q_value" is the Benjamini-Hochberg false discovery rate-adjusted $p$-value across all 160 residue-position tests. Rows with background frequency of 0 and observed count of 0 are structural zeros; fold-enrichment and $p$ and $q$-values are reported as not applicable.

| position | amino_acid | observed_count | background_frequency | fold_enrichment | p_value | q_value | expected_count |
|---|---|---|---|---|---|---|---|
| X-2 | A | 76 | 0.13450292 | 0.75946704 | 0.99682329 | 1 | 100.070175 |
| X-2 | C | 0 | 0 | | | | 0 |
| X-2 | D | 11 | 0.02923977 | 0.50564516 | 0.99630735 | 1 | 21.754386 |
| X-2 | E | 96 | 0.13450292 | 0.95932679 | 0.6847581 | 1 | 100.070175 |
| X-2 | F | 4 | 0.02339181 | 0.22983871 | 0.99997489 | 1 | 17.4035088 |
| X-2 | G | 37 | 0.04385965 | 1.13387097 | 0.2398275 | 0.714153 | 32.631579 |
| X-2 | H | 18 | 0.02046784 | 1.18202765 | 0.2691432 | 0.76734444 | 15.2280702 |
| X-2 | I | 22 | 0.02923977 | 1.01129032 | 0.5083976 | 1 | 21.754386 |
| X-2 | K | 22 | 0.03508772 | 0.84274194 | 0.81953706 | 1 | 26.1052632 |
| X-2 | L | 24 | 0.05847953 | 0.5516129 | 0.99964424 | 1 | 43.5087719 |
| X-2 | M | 25 | 0.01461988 | 2.2983871 | 0.00014743 | 0.00282221 | 10.877193 |
| X-2 | N | 16 | 0.02046784 | 1.05069124 | 0.45549841 | 1 | 15.2280702 |
| X-2 | P | 249 | 0.27777778 | 1.20483871 | 0.00038121 | 0.00567574 | 206.666667 |
| X-2 | Q | 21 | 0.02631579 | 1.07258065 | 0.40305366 | 1 | 19.5789474 |
| X-2 | R | 13 | 0.02339181 | 0.74697581 | 0.88695004 | 1 | 17.4035088 |
| X-2 | S | 65 | 0.07894737 | 1.10663082 | 0.21445611 | 0.68258677 | 58.7368421 |
| X-2 | T | 16 | 0.01754386 | 1.22580645 | 0.23952359 | 0.714153 | 13.0526316 |
| X-2 | V | 24 | 0.02631579 | 1.22580645 | 0.18240105 | 0.6432037 | 19.5789474 |
| X-2 | W | 0 | 0 | | | | 0 |
| X-2 | Y | 5 | 0.00584795 | 1.14919355 | 0.4395752 | 1 | 4.35087719 |
| Y-2 | A | 67 | 0.12573099 | 0.71624156 | 0.99908893 | 1 | 93.5438597 |
| Y-2 | C | 0 | 0 | | | | 0 |
| Y-2 | D | 26 | 0.04678363 | 0.74697581 | 0.95209635 | 1 | 34.8070175 |
| Y-2 | E | 8 | 0.00584795 | 1.83870968 | 0.07422764 | 0.35523227 | 4.35087719 |
| Y-2 | F | 0 | 0 | | | | 0 |
| Y-2 | G | 14 | 0.01461988 | 1.28709677 | 0.20604695 | 0.67342173 | 10.877193 |
| Y-2 | H | 0 | 0 | | | | 0 |
| Y-2 | I | 17 | 0.01169591 | 1.95362903 | 0.00784107 | 0.07004691 | 8.70175439 |
| Y-2 | K | 58 | 0.07602339 | 1.02543424 | 0.44076304 | 1 | 56.5614035 |
| Y-2 | L | 8 | 0.00584795 | 1.83870968 | 0.07422764 | 0.35523227 | 4.35087719 |
| Y-2 | M | 18 | 0.01169591 | 2.06854839 | 0.00359753 | 0.04228289 | 8.70175439 |
| Y-2 | N | 20 | 0.04678363 | 0.57459677 | 0.99787999 | 1 | 34.8070175 |
| Y-2 | P | 315 | 0.41520468 | 1.01970695 | 0.33818022 | 0.93279109 | 308.912281 |
| Y-2 | Q | 25 | 0.04385965 | 0.76612903 | 0.93211892 | 1 | 32.631579 |
| Y-2 | R | 105 | 0.11695906 | 1.20665323 | 0.02529004 | 0.14734196 | 87.0175439 |
| Y-2 | S | 46 | 0.04385965 | 1.40967742 | 0.01373632 | 0.09203332 | 32.631579 |
| Y-2 | T | 14 | 0.02631579 | 0.71505376 | 0.92430623 | 1 | 19.5789474 |
| Y-2 | V | 3 | 0.00877193 | 0.45967742 | 0.95841082 | 1 | 6.52631579 |
| Y-2 | W | 0 | 0 | | | | 0 |
| Y-2 | Y | 0 | 0 | | | | 0 |
| X-1 | A | 87 | 0.13702624 | 0.85338023 | 0.95264622 | 1 | 101.947522 |
| X-1 | C | 0 | 0 | | | | 0 |
| X-1 | D | 18 | 0.02915452 | 0.82983871 | 0.81823534 | 1 | 21.6909621 |
| X-1 | E | 103 | 0.13411079 | 1.03228728 | 0.38028049 | 0.99916834 | 99.7784257 |
| X-1 | F | 22 | 0.02332362 | 1.26780914 | 0.15648101 | 0.58245707 | 17.3527697 |
| X-1 | G | 23 | 0.04373178 | 0.70689964 | 0.96941721 | 1 | 32.5364432 |
| X-1 | H | 24 | 0.02040816 | 1.58064516 | 0.02086403 | 0.13313235 | 15.1836735 |
| X-1 | I | 27 | 0.02915452 | 1.24475807 | 0.14763398 | 0.56741544 | 21.6909621 |

| | | | | | | | |
|---|---|---|---|---|---|---|---|
| X-1 | K | 17 | 0.03498542 | 0.6531138 | 0.97732035 | 1 | 26.0291545 |
| X-1 | L | 19 | 0.05830904 | 0.43797043 | 0.99999301 | 1 | 43.3819242 |
| X-1 | M | 22 | 0.01457726 | 2.02849462 | 0.0017516 | 0.02347148 | 10.8454811 |
| X-1 | N | 36 | 0.02040816 | 2.37096774 | 2.89E-06 | 9.69E-05 | 15.1836735 |
| X-1 | P | 256 | 0.27696793 | 1.24233164 | 3.63E-05 | 0.00097172 | 206.06414 |
| X-1 | Q | 19 | 0.02623907 | 0.97326762 | 0.57910604 | 1 | 19.5218659 |
| X-1 | R | 14 | 0.02332362 | 0.80678763 | 0.82449693 | 1 | 17.3527697 |
| X-1 | S | 45 | 0.0787172 | 0.76836918 | 0.97575866 | 1 | 58.5655977 |
| X-1 | T | 5 | 0.01749271 | 0.38418459 | 0.99650557 | 1 | 13.0145773 |
| X-1 | V | 6 | 0.02623907 | 0.30734767 | 0.99991069 | 1 | 19.5218659 |
| X-1 | W | 0 | 0 | | | | 0 |
| X-1 | Y | 1 | 0.0058309 | 0.23051075 | 0.98710466 | 1 | 4.33819242 |
| Y-1 | A | 54 | 0.12536443 | 0.57895724 | 0.99999884 | 1 | 93.271137 |
| Y-1 | C | 0 | 0 | | | | 0 |
| Y-1 | D | 29 | 0.04664723 | 0.83560148 | 0.86096005 | 1 | 34.7055394 |
| Y-1 | E | 7 | 0.0058309 | 1.61357527 | 0.14820553 | 0.56741544 | 4.33819242 |
| Y-1 | F | 0 | 0 | | | | 0 |
| Y-1 | G | 3 | 0.01457726 | 0.2766129 | 0.99869541 | 1 | 10.8454811 |
| Y-1 | H | 0 | 0 | | | | 0 |
| Y-1 | I | 13 | 0.01166181 | 1.49831989 | 0.10069759 | 0.4652923 | 8.67638484 |
| Y-1 | K | 77 | 0.07580175 | 1.36533292 | 0.00378653 | 0.04228289 | 56.3965015 |
| Y-1 | L | 3 | 0.0058309 | 0.69153226 | 0.808227 | 1 | 4.33819242 |
| Y-1 | M | 4 | 0.01166181 | 0.46102151 | 0.9739745 | 1 | 8.67638484 |
| Y-1 | N | 13 | 0.04664723 | 0.37457997 | 0.9999944 | 1 | 34.7055394 |
| Y-1 | P | 342 | 0.41690962 | 1.1025829 | 0.0101447 | 0.0781297 | 310.180758 |
| Y-1 | Q | 39 | 0.04373178 | 1.19865591 | 0.14306451 | 0.56741544 | 32.5364432 |
| Y-1 | R | 108 | 0.11661808 | 1.24475807 | 0.01049504 | 0.0781297 | 86.7638484 |
| Y-1 | S | 17 | 0.04373178 | 0.52249104 | 0.99914209 | 1 | 32.5364432 |
| Y-1 | T | 31 | 0.02623907 | 1.58796296 | 0.00905491 | 0.07583486 | 19.5218659 |
| Y-1 | V | 4 | 0.00874636 | 0.61469534 | 0.88970423 | 1 | 6.50728863 |
| Y-1 | W | 0 | 0 | | | | 0 |
| Y-1 | Y | 0 | 0 | | | | 0 |
| X1 | A | 99 | 0.13702624 | 0.97108785 | 0.6391926 | 1 | 101.947522 |
| X1 | C | 0 | 0 | | | | 0 |
| X1 | D | 27 | 0.02915452 | 1.24475807 | 0.14763398 | 0.56741544 | 21.6909621 |
| X1 | E | 73 | 0.13411079 | 0.73162109 | 0.99885536 | 1 | 99.7784257 |
| X1 | F | 9 | 0.02332362 | 0.51864919 | 0.99034647 | 1 | 17.3527697 |
| X1 | G | 17 | 0.04373178 | 0.52249104 | 0.99914209 | 1 | 32.5364432 |
| X1 | H | 12 | 0.02040816 | 0.79032258 | 0.82984197 | 1 | 15.1836735 |
| X1 | I | 15 | 0.02915452 | 0.69153226 | 0.94829409 | 1 | 21.6909621 |
| X1 | K | 20 | 0.03498542 | 0.76836918 | 0.90807355 | 1 | 26.0291545 |
| X1 | L | 51 | 0.05830904 | 1.17560484 | 0.13364243 | 0.56741544 | 43.3819242 |
| X1 | M | 12 | 0.01457726 | 1.10645161 | 0.40215582 | 1 | 10.8454811 |
| X1 | N | 13 | 0.02040816 | 0.8561828 | 0.74994795 | 1 | 15.1836735 |
| X1 | P | 301 | 0.27988338 | 1.44549451 | 1.78E-13 | 2.39E-11 | 208.233236 |
| X1 | Q | 14 | 0.02623907 | 0.71714456 | 0.92252169 | 1 | 19.5218659 |
| X1 | R | 14 | 0.02332362 | 0.80678763 | 0.82449693 | 1 | 17.3527697 |
| X1 | S | 48 | 0.0787172 | 0.81959379 | 0.93746224 | 1 | 58.5655977 |
| X1 | T | 6 | 0.01749271 | 0.46102151 | 0.98984882 | 1 | 13.0145773 |
| X1 | V | 10 | 0.02332362 | 0.57627688 | 0.97937622 | 1 | 17.3527697 |
| X1 | W | 0 | 0 | | | | 0 |
| X1 | Y | 3 | 0.0058309 | 0.69153226 | 0.808227 | 1 | 4.33819242 |
| Y1 | A | 94 | 0.12536443 | 1.00781445 | 0.4843782 | 1 | 93.271137 |
| Y1 | C | 0 | 0.00291545 | 0 | 1 | 1 | 2.16909621 |
| Y1 | D | 19 | 0.04664723 | 0.54746304 | 0.99886704 | 1 | 34.7055394 |
| Y1 | E | 11 | 0.0058309 | 2.53561828 | 0.00500198 | 0.04787606 | 4.33819242 |
| Y1 | F | 0 | 0 | | | | 0 |
| Y1 | G | 2 | 0.01166181 | 0.23051075 | 0.99841498 | 1 | 8.67638484 |
| Y1 | H | 0 | 0 | | | | 0 |
| Y1 | I | 11 | 0.01166181 | 1.26780914 | 0.25551501 | 0.74432634 | 8.67638484 |
| Y1 | K | 59 | 0.07580175 | 1.04616419 | 0.37858053 | 0.99916834 | 56.3965015 |
| Y1 | L | 9 | 0.0058309 | 2.07459677 | 0.03279126 | 0.18308452 | 4.33819242 |
| Y1 | M | 3 | 0.01166181 | 0.34576613 | 0.9921776 | 1 | 8.67638484 |

| | | | | | | | |
|---|---|---|---|---|---|---|---|
| Y1 | N | 19 | 0.04664723 | 0.54746304 | 0.99886704 | 1 | 34.7055394 |
| Y1 | P | 306 | 0.41690962 | 0.98652154 | 0.63540478 | 1 | 310.180758 |
| Y1 | Q | 42 | 0.04373178 | 1.29086022 | 0.05823403 | 0.30256878 | 32.5364432 |
| Y1 | R | 94 | 0.11661808 | 1.08340054 | 0.21903904 | 0.68258677 | 86.7638484 |
| Y1 | S | 40 | 0.04373178 | 1.22939068 | 0.10828291 | 0.48366365 | 32.5364432 |
| Y1 | T | 29 | 0.02623907 | 1.48551374 | 0.02480773 | 0.14734196 | 19.5218659 |
| Y1 | V | 6 | 0.00874636 | 0.92204301 | 0.63297927 | 1 | 6.50728863 |
| Y1 | W | 0 | 0 | | | | 0 |
| Y1 | Y | 0 | 0 | | | | 0 |
| X2 | A | 84 | 0.1374269 | 0.82155113 | 0.97925175 | 1 | 102.245614 |
| X2 | C | 0 | 0 | | | | 0 |
| X2 | D | 12 | 0.02923977 | 0.5516129 | 0.99201513 | 1 | 21.754386 |
| X2 | E | 84 | 0.13450292 | 0.83941094 | 0.96498438 | 1 | 100.070175 |
| X2 | F | 7 | 0.02339181 | 0.40221774 | 0.99856863 | 1 | 17.4035088 |
| X2 | G | 27 | 0.04385965 | 0.82741936 | 0.86543198 | 1 | 32.631579 |
| X2 | H | 16 | 0.02046784 | 1.05069124 | 0.45549841 | 1 | 15.2280702 |
| X2 | I | 41 | 0.02923977 | 1.88467742 | 0.00011537 | 0.00257666 | 21.754386 |
| X2 | K | 18 | 0.03508772 | 0.68951613 | 0.96307873 | 1 | 26.1052632 |
| X2 | L | 47 | 0.05555556 | 1.13709677 | 0.20192432 | 0.67342173 | 41.3333333 |
| X2 | M | 4 | 0.01461988 | 0.36774194 | 0.99483892 | 1 | 10.877193 |
| X2 | N | 25 | 0.02046784 | 1.64170507 | 0.01234722 | 0.08708041 | 15.2280702 |
| X2 | P | 209 | 0.28070175 | 1.00075605 | 0.50875376 | 1 | 208.842105 |
| X2 | Q | 19 | 0.02631579 | 0.97043011 | 0.58418586 | 1 | 19.5789474 |
| X2 | R | 11 | 0.02339181 | 0.63205645 | 0.9610971 | 1 | 17.4035088 |
| X2 | S | 79 | 0.07894737 | 1.34498208 | 0.00488388 | 0.04787606 | 58.7368421 |
| X2 | T | 42 | 0.01754386 | 3.21774194 | 8.94E-11 | 5.99E-09 | 13.0526316 |
| X2 | V | 16 | 0.02339181 | 0.91935484 | 0.66666278 | 1 | 17.4035088 |
| X2 | W | 0 | 0 | | | | 0 |
| X2 | Y | 3 | 0.00584795 | 0.68951613 | 0.80978182 | 1 | 4.35087719 |
| Y2 | A | 90 | 0.12280702 | 0.98502304 | 0.57737804 | 1 | 91.3684211 |
| Y2 | C | 1 | 0.00292398 | 0.45967742 | 0.88680296 | 1 | 2.1754386 |
| Y2 | D | 40 | 0.04678363 | 1.14919355 | 0.2049859 | 0.67342173 | 34.8070175 |
| Y2 | E | 14 | 0.00584795 | 3.21774194 | 0.00016953 | 0.00283956 | 4.35087719 |
| Y2 | F | 0 | 0 | | | | 0 |
| Y2 | G | 12 | 0.01169591 | 1.37903226 | 0.16782712 | 0.60780634 | 8.70175439 |
| Y2 | H | 0 | 0 | | | | 0 |
| Y2 | I | 14 | 0.01169591 | 1.60887097 | 0.05870738 | 0.30256878 | 8.70175439 |
| Y2 | K | 58 | 0.07602339 | 1.02543424 | 0.44076304 | 1 | 56.5614035 |
| Y2 | L | 5 | 0.00584795 | 1.14919355 | 0.4395752 | 1 | 4.35087719 |
| Y2 | M | 3 | 0.01169591 | 0.34475807 | 0.99233571 | 1 | 8.70175439 |
| Y2 | N | 22 | 0.04678363 | 0.63205645 | 0.99286753 | 1 | 34.8070175 |
| Y2 | P | 298 | 0.41812866 | 0.95792917 | 0.84374741 | 1 | 311.087719 |
| Y2 | Q | 23 | 0.04385965 | 0.70483871 | 0.97051759 | 1 | 32.631579 |
| Y2 | R | 91 | 0.11695906 | 1.04576613 | 0.34109525 | 0.93279109 | 87.0175439 |
| Y2 | S | 62 | 0.04385965 | 1.9 | 1.66E-06 | 7.40E-05 | 32.631579 |
| Y2 | T | 4 | 0.02631579 | 0.20430108 | 0.9999962 | 1 | 19.5789474 |
| Y2 | V | 7 | 0.00877193 | 1.07258065 | 0.4779806 | 1 | 6.52631579 |
| Y2 | W | 0 | 0 | | | | 0 |
| Y2 | Y | 0 | 0 | | | | 0 |

**Table S3. Summary of classical interchain hydrogen-bond counts from molecular dynamics simulations of the CMPs.**

For each peptide, the table reports the number of analyzed frames (*n*_frames) and summary statistics for the number of classical interchain hydrogen bonds per frame across the 500 ns trajectory, including hbond_mean, hbond_median, hbond_sd, hbond_min, and hbond_max. Peptides are additionally annotated by variant class, primary group, and subgroup used for the comparisons in Figure 5B.

| peptide | variant | n_frames | hbond_mean | hbond_median | hbond_sd | hbond_min | hbond_max | group | subgroup |
|---|---|---|---|---|---|---|---|---|---|
| A1-G382 | WT | 1250 | 33.5536 | 34 | 2.73823399 | 12 | 37 | WT | WT |
| A1-G382R | G>R | 1250 | 28.8096 | 29 | 3.58592177 | 9 | 37 | G>R | G>R |
| A1-G382S | G>S | 1250 | 31.772 | 32 | 2.6570781 | 11 | 37 | G>S | G>S nonGP |
| A1-G448 | WT | 1250 | 31.5496 | 32 | 2.84190094 | 15 | 39 | WT | WT |
| A1-G448S | G>S | 1250 | 30.1208 | 31 | 4.25590149 | 8 | 37 | G>S | G>S nonGP |
| A1-G661 | WT | 1250 | 33.024 | 34 | 4.38091063 | 11 | 37 | WT | WT |
| A1-G661S | G>S | 1250 | 28.5064 | 30 | 5.0111524 | 6 | 36 | G>S | G>S GP |
| A1-G832 | WT | 1250 | 32.8656 | 33 | 2.6512563 | 14 | 38 | WT | WT |
| A1-G832S | G>S | 1250 | 30.0344 | 31 | 3.91211712 | 11 | 37 | G>S | G>S nonGP |
| A1-G862 | WT | 1250 | 32.7496 | 34 | 4.05571929 | 10 | 38 | WT | WT |
| A1-G862S | G>S | 1250 | 29.6024 | 30 | 3.64160999 | 10 | 36 | G>S | G>S GP |
| A1-G964 | WT | 1250 | 33.584 | 34 | 3.15285528 | 9 | 37 | WT | WT |
| A1-G964S | G>S | 1250 | 28.4888 | 29 | 3.89867083 | 11 | 35 | G>S | G>S GP |
| A2-G661 | WT | 1250 | 32.484 | 34 | 5.62065988 | 3 | 37 | WT | WT |
| A2-G661S | G>S | 1250 | 30.8904 | 31 | 2.67389372 | 11 | 36 | G>S | G>S nonGP |
| T3-G16 | WT | 1250 | 32.2464 | 33 | 3.52505714 | 12 | 38 | WT | WT |
| T3-G16S | G>S | 1250 | 28.012 | 29 | 4.44887514 | 10 | 34 | G>S | G>S GP |
| T3-G19S | G>S | 1250 | 30.62 | 31 | 3.36497525 | 9 | 37 | G>S | G>S nonGP |
| T3-G277 | WT | 1250 | 32.1616 | 33 | 4.25689821 | 12 | 38 | WT | WT |
| T3-G277R | G>R | 1250 | 29.3784 | 30 | 3.76852423 | 9 | 36 | G>R | G>R |
| T3-G415 | WT | 1250 | 32.8976 | 33 | 2.67642389 | 13 | 38 | WT | WT |
| T3-G415R | G>R | 1250 | 28.4736 | 29 | 3.38858216 | 11 | 36 | G>R | G>R |
| T3-G415S | G>S | 1250 | 28.8848 | 29 | 2.92656467 | 12 | 36 | G>S | G>S GP |
| T3-G619 | WT | 1250 | 32.1584 | 33 | 4.162111 | 9 | 37 | WT | WT |
| T3-G619R | G>R | 1250 | 27.4672 | 28 | 3.40126139 | 7 | 34 | G>R | G>R |
| T3-G70 | WT | 1250 | 31.7152 | 33 | 5.24565558 | 4 | 38 | WT | WT |
| T3-G70R | G>R | 1250 | 28.8864 | 29 | 2.03056485 | 14 | 35 | G>R | G>R |
